# Paired heavy and light chain signatures contribute to potent SARS-CoV-2 neutralization in public antibody responses

**DOI:** 10.1101/2020.12.31.424987

**Authors:** Bailey B. Banach, Gabriele Cerutti, Ahmed S. Fahad, Chen-Hsiang Shen, Matheus Oliveira de Souza, Phinikoula S. Katsamba, Yaroslav Tsybovsky, Pengfei Wang, Manoj S. Nair, Yaoxing Huang, Irene M. Francino Urdániz, Paul J. Steiner, Matias Gutiérrez-González, Lihong Liu, Sheila N. López Acevedo, Alexandra Nazzari, Jacy R. Wolfe, Yang Luo, Adam S. Olia, I-Ting Teng, Jian Yu, Tongqing Zhou, Eswar R. Reddem, Jude Bimela, Xiaoli Pan, Bharat Madan, Amy D. Laflin, Rajani Nimrania, Kwon-Tung Yuen, Timothy A. Whitehead, David D. Ho, Peter D. Kwong, Lawrence Shapiro, Brandon J. DeKosky

## Abstract

Understanding protective mechanisms of antibody recognition can inform vaccine and therapeutic strategies against SARS-CoV-2. We discovered a new antibody, 910-30, that targets the SARS-CoV-2 ACE2 receptor binding site as a member of a public antibody response encoded by IGHV3-53/IGHV3-66 genes. We performed sequence and structural analyses to explore how antibody features correlate with SARS-CoV-2 neutralization. Cryo-EM structures of 910-30 bound to the SARS-CoV-2 spike trimer revealed its binding interactions and ability to disassemble spike. Despite heavy chain sequence similarity, biophysical analyses of IGHV3-53/3-66 antibodies highlighted the importance of native heavy:light pairings for ACE2 binding competition and for SARS-CoV-2 neutralization. We defined paired heavy:light sequence signatures and determined antibody precursor prevalence to be ~1 in 44,000 human B cells, consistent with public antibody identification in several convalescent COVID-19 patients. These data reveal key structural and functional neutralization features in the IGHV3-53/3-66 public antibody class to accelerate antibody-based medical interventions against SARS-CoV-2.

**Highlights:**

- A molecular study of IGHV3-53/3-66 public antibody responses reveals critical heavy and light chain features for potent neutralization
- Cryo-EM analyses detail the structure of a novel public antibody class member, antibody 910-30, in complex with SARS-CoV-2 spike trimer
- Cryo-EM data reveal that 910-30 can both bind assembled trimer and can disassemble the SARS-CoV-2 spike
- Sequence-structure-function signatures defined for IGHV3-53/3-66 class antibodies including both heavy and light chains
- IGHV3-53/3-66 class precursors have a prevalence of 1:44,000 B cells in healthy human antibody repertoires

## Introduction

SARS-CoV-2 emerged in late 2019 into human populations, causing coronavirus disease 2019 (COVID-19) with complications including respiratory and cardiac failure in severe cases (Gorbalenya and et al., 2020; Guan et al., 2020; Jiang et al., 2020; Zhou et al., 2020a; Zho et al., 2020). The highly infectious nature of SARS-CoV-2, significant prevalence of severe disease, and widespread transmission by asymptomatic and pre-symptomatic individuals has led to immense global, social, and economic disruption (Cucinotta and Vanelli, 2020; Liu et al., 2020b). SARS-CoV-2 marks the third known emergence of a novel beta-coronavirus in the past two decades, following its closest documented human pathogen severe acute respiratory syndrome coronavirus (SARS-CoV) in 2002, and the next closest, Middle East respiratory syndrome coronavirus (MERS-CoV) in 2012 (Cui et al., 2019; Gorbalenya and et al., 2020; Graham and Baric, 2010; Ksiazek et al., 2003; de Wit et al., 2016; Zaki et al., 2012). Both SARS and SARS-CoV-2 infect human cells by binding to the angiotensin convertase II receptor (ACE2) via the trimeric spike (S) class I fusion protein (Hoffmann et al., 2020; Wrapp et al., 2020a). The S protein comprises two subunits, S1 and S2. The S1 subunit contains a receptor binding domain (RBD), which binds to ACE2. To enter cells, S must undergo a protease cleavage event that allows S1 to shed and expose the hydrophobic fusion peptide of the S2 subunit. SARS coronavirus predominantly enters cells via endosomes, assisted by cathepsin cleavage in the low pH (5.5-4.5) endosomal environment. SARS-CoV-2 acquired a new protease cleavage site that enables entry either at the cell surface after cleavage with TMPRSS2, or inside endosomes via protease cleavage similar to SARS, and the route of SARS-CoV-2 entry is likely dependent on the protease expression profile in target cells (Ou et al., 2020; Tang et al., 2020). ACE2 interactions appear to play a role in the pre-fusion shedding of S1 (Benton et al., 2020; Cai et al., 2020).

A detailed understanding of SARS-CoV-2 molecular vulnerabilities to antibody neutralization can accelerate progress in medical interventions such as antibody drug therapies and vaccines. Antibodies from several COVID-19 patients have revealed the presence of public antibody responses that target SARS-CoV-2 via shared genetic elements and structural recognition modes in the IGHV3-53 and IGHV3-66 heavy chain V-genes. Members of this public antibody class target a conserved epitope on RBD on the S1 subunit that overlaps with the ACE2 binding site (Barnes et al., 2020; Brouwer et al., 2020; Cao et al., 2020; Chi et al., 2020; Du et al., 2020; Hansen et al., 2020; Hurlburt et al., 2020; Liu et al., 2020a; Rogers et al., 2020; Seydoux et al., 2020; Shi et al., 2020; Wu et al., 2020b; Yuan et al., 2020a). IGHV3-53/3-66 public class antibodies share common genetic features including IGHV-gene-encoded motifs NY in the CDR-H1, SGGS in the CDR-H2, a relatively short CDR-H3 length, and comparatively low levels of antibody somatic hypermutation (Barnes et al., 2020; Du et al., 2020; Wu et al., 2020a; Yuan et al., 2020a). Preliminary analysis of class light chain features show the inclusion of both kappa and lambda light chains in antibodies of this class (Catalan-Dibene, 2020; Du et al., 2020; Wang et al., 2020a; Wrapp et al., 2020b; Wu et al., 2020b, 2020b; Yuan et al., 2020a). Despite strong similarities in heavy chain gene signatures, IGHV3-53/3-66 anti-RBD antibodies show a broad range of neutralization potencies (IC50’s from 0.003 to 2.547 μg/mL), (Brouwer et al., 2020; Cao et al., 2020; Ju et al., 2020; Liu et al., 2020a; Robbiani et al., 2020; Rogers et al., 2020; Shi et al., 2020; Wu et al., 2020b; Yuan et al., 2020; Zost et al., 2020). Given the low somatic hypermutation observed and the importance of germline-encoded recognition motifs, it remains unclear what unique molecular features lead to the diverse range of SARS-CoV-2 neutralization potencies among IGHV3-53/3-66 class members.

SARS-CoV-2 S displays a pH-dependent conformational switch that causes the ‘up’ position of the RBD to rotate to a ‘down’ position (Walls et al., 2020; Zhou et al., 2020c). The RBD ‘up’ position is required for ACE2 engagement, as well as antibody binding for the IGHV3-53/3-66 class (Du et al., 2020; Walls et al., 2020; Wrapp et al., 2020b). A mutational variant recently emerged that influences the RBD “up” vs. “down” state (D614G) that now constitutes >97% of isolates world-wide (Korber et al., 2020; Long et al., 2020; Volz et al., 2020; Yurkovetskiy et al., 2020; Zhang et al., 2020). The D614G mutation is proximal to the RBD in the spike structure, and D614G appears to favor more RBD ‘up’ at both serological and endosomal pH (Grubaugh et al., 2020; Yurkovetskiy et al., 2020; Zhou et al., 2020c). The D614G substitution enhances viral infectivity, competitive fitness, and transmission, and may have important implications for antibody-based and vaccine interventions (Hou et al., 2020; Mansbach et al., 2020; Yurkovetskiy et al., 2020; Zhang et al., 2020); further investigations into the effects of D614G on IGHV3-53/3-66 class recognition and neutralization are required (Grubaugh et al., 2020).

Here we discovered a new member of the IGHV3-53/3-66 antibody class, mAb 910-30, with moderate neutralization capacity. To better understand the features of potent IGHV3-53/3-66 class neutralization, we explored molecular and genetic features of 910-30 and other related antibodies, including heavy and light chain structural recognition motifs, biophysical correlates of neutralization, and the influence of D614 vs. D614G variants on IGHV3-53/3-66 class member interactions. Our study provides a detailed molecular understanding of how the public IGHV3-53/3-66 class leverages native heavy and light chain binding contributions against SARS-CoV-2, providing important data to accelerate medical interventions against SARS-CoV-2’s vulnerable RBD epitope.

## Results

### Isolation and structural characterization of a novel neutralizing class member

We screened the immune repertoire of a COVID-19 convalescent patient, Donor 910 (To et al., 2020) treated at Hong Kong University, to identify a new member of the public IGHV3-53/3-66 antibody class. ELISA assays of Donor 910 serum showed potent S trimer recognition, and pseudovirus neutralization assays confirmed that Donor 910 serum showed high SARS-CoV-2 neutralization titers (**Fig. S1**) (Wang et al., 2020b). Based on these data, Donor 910 cryopreserved peripheral blood mononuclear cells (PBMCs) were selected for analysis using a recently described method to clone natively paired heavy and light chain antibody genes into yeast Fab display for functional screening (Wang et al., 2018). Yeast antibody display libraries were screened for binding against two different SARS-CoV-2 S protein probes, a His-labeled S-Trimer and a biotinylated S-Trimer, by fluorescence-activated cell sorting (FACS) (**Fig. S1**). Bioinformatic interrogation of yeast display NGS data revealed that one monoclonal antibody (mAb 910-30) enriched 90-fold in the round 2 sorted library against Biotin-22 S protein, and 2,296-fold in round 3 sorted libraries against His-labeled S protein. Based on these strong enrichment signals, mAb 910-30 was expressed as IgG in HEK293 cells for neutralization assays. 910-30 showed a half-maximal inhibitory concentrations (IC_50_) against a VSV SARS-CoV-2 pseudovirus (Liu et al., 2020a) of 0.071 μg/mL, and 0.142 μg/mL against authentic SARS-CoV-2 (**Fig. 1A**) shown in comparison to the previously reported mAb CR3022 (Huo et al., 2020; Ter Meulen et al., 2006; Yuan et al., 2020b).

**Fig. 1.**
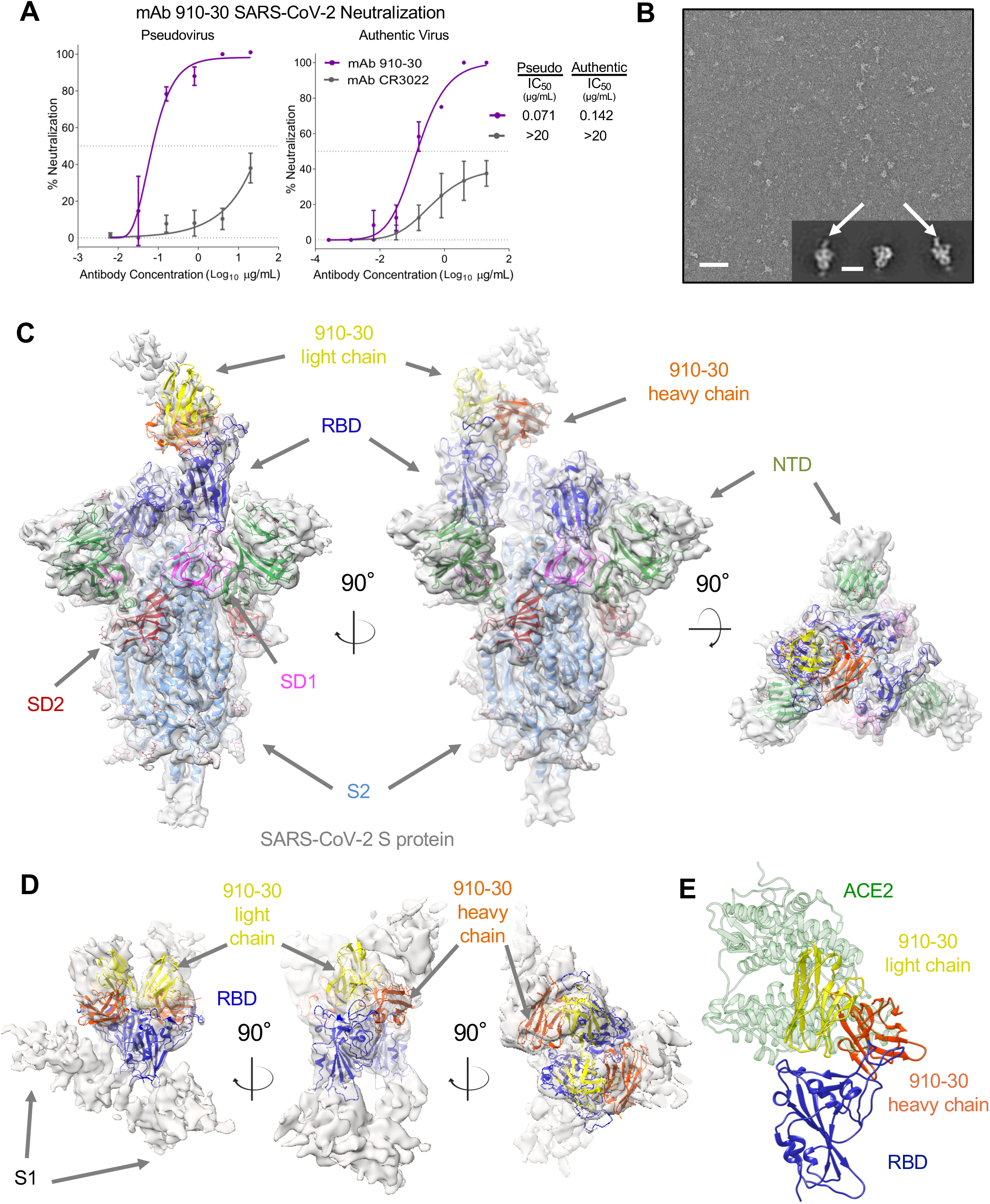
A novel SARS-CoV-2 neutralizer in the reproducible IGHV3-53/3-66 antibody class targets the ACE2 binding site of both ordered and disassembled spike. **(A)** The novel SARS-CoV-2 neutralizing antibody 910-30 shows moderately potent neutralization capacity compared to the control mAb CR3022 in both VSV-pseudo-type virus and authentic virus assays. **(B)** Negative-staining electron microscopy at pH 5.5 revealed 910-30 Fab bound to SARS-CoV-2 S2P protein. A representative micrograph is shown. Inset shows representative 2D class averages; arrows point to bound Fab fragments. Scale bars: 50 nm (micrographs) 20 nm (2D class averages). **(C)** Cryo-EM map and molecular model of 910-30 Fab in complex with SARS-CoV-2 spike at 4.75 Å resolution. Only one conformation, with 1 Fab bound to 1 RBD up, is observed when mixing Fab and spike in a 1:1 molar ratio at pH 5.5. NTD is colored in green, RBD in blue, SD1 in magenta, SD2 in red, S2 in light blue, 910-30 antibody heavy chain in orange, 910-30 light chain in yellow. **(D)** Cryo-EM map obtained when 910-30 Fab and spike are mixed in a 9:1 molar ratio at pH 5.5. The only observed species is a mostly disordered spike in which RBD and 91030 Fab still fit the map consistently with the properly folded spike:910-30 complex shown in (C). RBD is shown in blue, 910-30 heavy chain in orange, 910-30 light chain in yellow. **(E)** The structural superposition of ACE2 (PDB entry 6M0J) and 910-30 (PDB entry 7KS9) in complex with SARS-CoV-2 RBD shows a representative ACE2 competition mechanism defining IGHV3-53/3-66 class neutralization. ACE2 is colored in green, 910-30 heavy chain in orange, 910-30 light chain in yellow, RBD in blue.

Next we characterized 910-30 structural recognition by cryo-electron microscopy (cryo-EM). Negative-staining electron microscopy revealed particles of 910-30 Fab bound to SARS-CoV-2 S2P at pH 5.5 (**Fig. 1B**) (Zhou et al., 2020c), and also across a broader pH range of 4.0-7.4 (**Fig. S1**). Subsequent Cryo-EM mapping and molecular modeling of 910-30 Fab in complex with SARS-CoV-2 S2P protein at pH 5.5 showed 1 Fab bound to 1 RBD in the up position when mixing Fab and spike in a 1:1 molar ratio (**Fig. 1C, Fig. S2, Supplemental Table 1**), whereas a 9:1 Fab:spike molar ratio revealed mostly disordered spike (**Fig. 1D**, **Fig. S2, Supplemental Table 1**), with an RBD that still fit the Cryo-EM map consistent with **Figure 1C**. Structural modeling of ACE2 (PDB entry 6M0J) and 910-30 (PDB entry 7KS9) in complex with SARS-CoV-2 RBD confirmed targeting of the ACE2 binding site (**Fig. 1E**). Analysis of soluble 910-30 IgG recognition of yeast-displayed aglycosylated N343Q RBD(333-537) confirmed that 910-30 recognizes a glycan-independent region (**Fig. S3A**) (Starr et al., 2020). Antibody titrations showed a 910-30 IgG K_D_ to RBD of 230 pM (191 - 268 pM 95% confidence interval) (**Fig. S3B**), and that 910-30 competes with human ACE2 (hACE2) for binding to RBD, consistent with IGHV3-53/3-66 class membership (**Fig. S3C**).

### Potent antibodies of the IGHV3-53/3-66 class compete strongly with ACE2 for binding to spike

Structural analysis of IGHV3-53/3-66 germline-encoded antibody recognition shows substantial overlap between the ACE2 binding site and the shared class epitope footprint (**Fig. 2A**) (Barnes et al., 2020; Cao et al., 2020; Hansen et al., 2020; Ju et al., 2020; Liu et al., 2020a; Shi et al., 2020; Walls et al., 2020; Wu et al., 2020b). Despite low reported somatic hypermutation and shared epitope targets, reported IGHV3-53/3-66 antibody class members show a broad range of neutralization potencies (**Fig. 2B, Supplemental Table 2**). To better understand molecular features of potent antibody neutralization for this class, we assessed biophysical performance of a small panel of weak, moderate, and potent IGHV3-53/3-66 public antibody class members. We selected the antibodies 1-20 (a potent neutralizer), the new 910-30 (a moderate neutralizer), and B38 (a weak neutralizer), along with a VH-gene matched control (mAb 4-3) that neutralizes poorly and likely targets a different site on RBD (**Supplemental Table 3**) (Wu et al., 2020a; Zhou et al., 2020a). Preliminary IgG ELISA analysis revealed that class members show similar binding to RBD, whereas the more potent class members (1-20, 910-30) bound more tightly to full-length spike (**Fig. 2C**). Pseudovirus & authentic virus neutralization show a range of two orders of magnitude in IC50 neutralization potencies for the selected panel (**Fig. 2D**), confirming that antibody neutralization within the class is driven by more complex parameters than simple recognition of the ACE2 binding site on RBD.

**Fig. 2.**
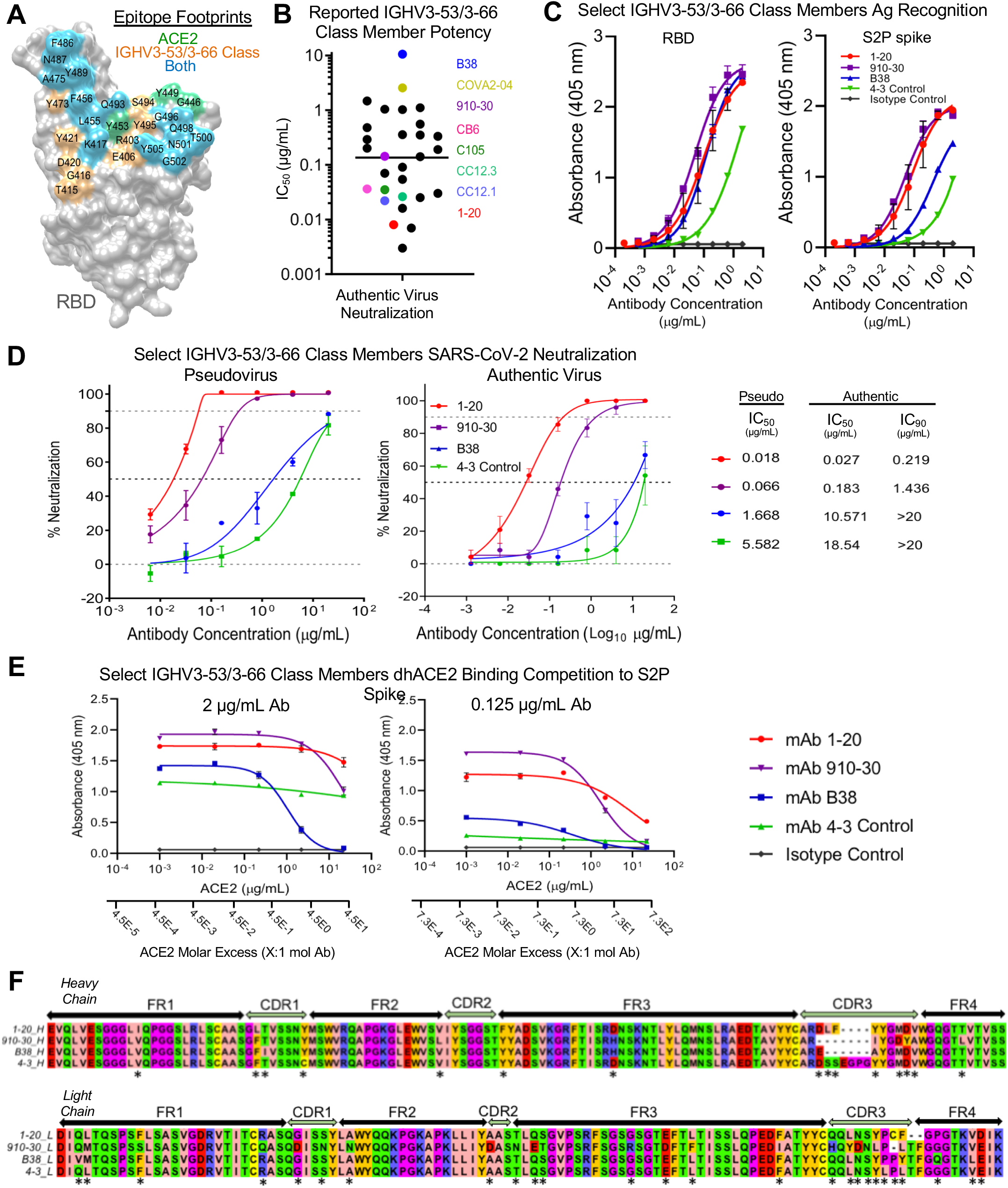
IGHV3-53/3-66 class neutralization potency is driven by strong competition with ACE2 for spike S2P recognition. **(A)** Epitope footprint comparison between ACE2 binding site and the IGHV3-53/3-66 class epitope on RBD show substantial overlap. Residues interacting with ACE2 only are shown in green, residues targeted by the IGHV3-53/3-66 class only are shown in orange, residues that overlap are shown in light blue. Other sites on RBD are represented in gray. **(B)** Dot-chart of reported wild-type authentic virus neutralization IC50 titers for previously published IGHV3-53/3-66 anti-RBD antibodies. Data were plotted without correcting for any differences in neutralization assay protocols. Line indicates the mean of IC50 values. A list of antibodies, IC50 values, and citations are provided in Supplemental Table 2. **(C)** IgG ELISA binding titrations for select IGHV3-53/3-66 class members against S2P spike and RBD antigens, with an IGHV gene-matched control (mAb 4-3) and an isotype control. **(D)** Pseudovirus & authentic virus neutralization show that IC50 neutralization potency ranges two orders of magnitude between the selected IGHV3-53/3-66 class members, along with an IGHV gene-matched control (mAb 4-3). **(E)** dhACE2 competition ELISA against SARS-CoV-2 S2P spike showing constant IgG concentrations with increasing dhACE2 (ACE2) concentrations. dhACE2 concentration is provided as both μg/mL and as ACE2 molar excess units. **(F)** Sequence alignment of heavy chain (upper) and light chain (lower) genes selected for detailed investigation. 4-3 is an anti-RBD antibody encoded by IGHV3-66 that does not compete with ACE2, and serves as a heavy chain gene-matched control.

Next we assessed the ability of potent and weak neutralizers to compete with dimerized human ACE2 (dhACE2) for binding to spike (S2P). In a competition ELISA using antibody pre-mixed with serial dilutions of dhACE2, we found that more potently neutralizing class members competed more strongly with dhACE2 compared to less potent Abs (**Fig. 2E**). The most potent mAb (1-20) required 73 dhACE2 molecules for 50% binding inhibition of 1 IgG molecule. 1-20 was 6-fold more competitive with dhACE2 than the moderate neutralizer 910-30 (dhACE2 molar excess IC50 = 12), and 150-fold more competitive than the weak neutralizer B38 (dhACE2 molar excess IC_50_ = 0.48). Sequence analysis revealed high similarity with low levels of somatic hypermutation, as previously reported for the IGHV3-53/3-66 antibody class (**Fig. 2F**) (Hurlburt et al., 2020; Yuan et al., 2020a). Given the broad variations in functional potency despite high sequence similarity, we next sought to understand the key contributions in heavy and light chain sequence signatures that lead to SARS-CoV-2 neutralization.

### Unique heavy and light chain interactions drive potent neutralization for the IGHV3-53/3-66 neutralizing antibody class

IGHV3-53/IGHV3-66 anti-SARS-CoV-2 antibodies show diverse light chain usage, with the two defining heavy chain genes (IGHV3-53 and IGHV3-66) pairing with at least 14 other light chain genes (**Supplemental Table 2**). To help understand the influence of light chain pairings, we constructed a panel of 12 non-native heavy:light swapped antibody variants from four IGHV3-53/3-66-encoded mAbs (1-20, 910-30, B38, and 4-3 included as an IGHV gene control). 11/12 non-native antibodies expressed successfully and were assayed for SARS-CoV-2 pseudovirus neutralization. Heavy:light swap data revealed a substantial loss in neutralization for nearly all non-native heavy:light combinations, with only the most potent antibody heavy chain (1-20) achieving significant neutralization with a non-native light chain (**Fig. 3A**). More potent neutralization of non-native heavy:light combinations was also correlated with strong dhACE2 competition (**Fig. 3B**), consistent with the natively paired heavy:light dhACE2 competition ELISA data (**Fig. 2E**). As all four heavy chain genes have low somatic hypermutation & high sequence similarity, we had not anticipated widespread loss of performance when swapping light chains among class members (**Fig. 2F, Supplemental Table 3**). In contrast, **Figure 3A** shows that native light chains make substantial contributions to neutralization performance, and that non-native heavy and light chain pairings show reduced antibody performance, despite the high degree of sequence similarity among heavy chains.

**Fig. 3.**
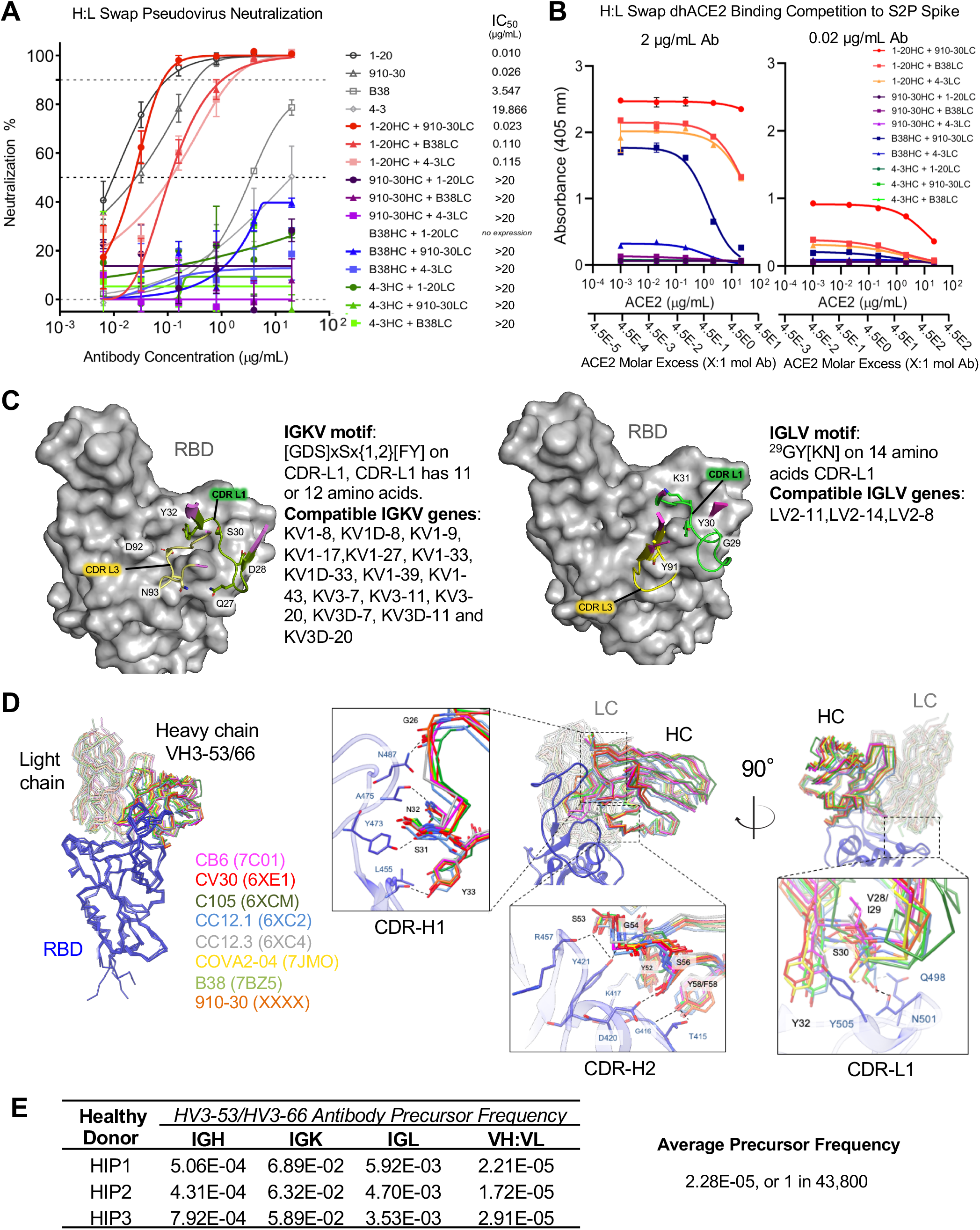
Heavy and light chain analyses reveal critical contributions of both VH and VL for potent antibody neutralization in the IGHV3-53/3-66 antibody class. **(A)** Heavy and light chain swap neutralization panel produced from four mAbs (1-20, 910-30, B38, and 4-3 included as an IGHV gene control) assayed for SARS-CoV-2 pseudovirus neutralization. **(B)** dhACE2 competition ELISA against SARS-CoV-2 S2P protein. Constant concentrations of heavy-light-swapped IgG were titrated with varying dhACE2 (ACE2) concentrations. dhACE2 concentration is provided in μg/mL, and also as ACE2 molar excess units. **(C)** Combined structure and sequences analyses reveal IGHV3-53/3-66 class light chain kappa (left panel) and lambda (right panel) genetic elements associated with the RBD contact interface. CDR-L1 residues are not specific to IGKV1-33 (910-30) and IGLV2-8 (C105) (Table S4). **(D)** Left panel: structural superposition of IGHV3-53/3-66 Fab variable domains in complex with RBD shows the same binding orientation for 8 different class antibodies aligned on RBD. RBD is shown in blue, Fab heavy chain in solid color, Fab light chain in transparent color, according to antibody name and the PDB code (shown in parentheses). Right panel: close-up views of the Fab:RBD interface for the eight IGHV3-53/3-66 antibodies superimposed on RBD. Conserved interactions of CDR-H1, −H2 and −L1 define the structural signatures responsible for viral neutralization by the IGHV3-53/3-66 antibody class. **(E)** Estimated probability of IGHV3-53/3-66 class pre-cursor antibodies derived from healthy donor (HIP1, HIP2, HIP3) immune repertories.

To better understand the paired heavy and light chain determinants of antibody recognition, we performed a structure-based alignment to analyze residue interactions and identify possible light chain signatures of class membership (Lefranc et al., 2003; Zhu et al., 2013). We followed numerous anti-RBD IGHV3-53/3-66 antibody lineages paired with different light chain V genes including: KV1-33, KV1-9, KV1-39, KV3-20, and LV2-8 (Barnes et al., 2020; Hurlburt et al., 2020; Shi et al., 2020; Wu et al., 2020b; Yuan et al., 2020a). Structural analyses of antibody contact sites revealed that conserved residues in both VH and VL genes contributed 56-75% of binding surface area (BSA) (**Fig. 3C and Fig. S3D**). We confirmed alignments with previously defined signatures as a control (**Fig. S3D**). **Supplemental Figure 3D** shows the IGHV3-53/3-66 heavy chain projected surface with its germline encoded amino acids, including the interaction residues ^31^SNY on CDR-H1 and ^52^YSGxSxY, where x indicates any residue, on CDR-H2 that provide multiple hydrogen bonds interactions with Thr415, Gly416, Lys417, Asp420, Tyr421, Leu455, Tyr473, Ala475, and Asn487 in the RBD. We verified previous reports that these class sequences have shorter CDR-H3 lengths of 6-11 amino acids (**Fig. S3D**), and we also note that those CDR-H1 and CDR-H2 motifs are only present in IGHV3-53/3-66 genes. Sequence-structure alignments also revealed that kappa chain class members use a conserved [DGS]xSx{1,2}[FY] motif of 11 or 12 amino acids starting at residue 27a or 28 in the CDR-L1 to form hydrogen bonds with RBD residues Gln498 and Asn501 (**Fig. 3C**). In contrast, lambda chain binding in the class uses a different ^29^GY[KN] motif with 14 amino acids in the CDR-L1 that interacts with RBD residues Gly502 and Tyr505 (**Fig. 3C**). Thus we defined here the ^27a/28^[DGS]xSx{1,2}[FY] motif of 11 or 12 amino acids in the CDR-L1 as a signature of kappa chain class members, and the ^29^GY[KN] motif on 14 amino acids CDR-L1 as a signature of lambda class members. We did not observe a conserved binding motif in the CDR-L3 that contacts RBD, and we also note that the CDR-L1 motifs defined here can be encoded by multiple light chain genes (**Fig. 3C, Table S4**).

Structural comparison of eight IGHV3-53/3-66 class members shows that Fab variable domains bind RBD with the same orientation, reflecting a conserved heavy chain recognition mode and defining the relevant conserved light chain residues for heavy:light pairing (**Fig. 3D**). Structural alignment also reveals a strongly conserved hydrogen bond network responsible for RBD recognition by CDR-H1. The backbone carbonyl group of Gly26HC interacts with the amide group of Asn487; the backbone CO group of Ser31HC contacts the hydroxyl group of Tyr473; the side chain amide group of Asn32_HC_ contacts the carbonyl group of Ala475; and the hydroxyl group of Tyr33_HC_ acts as hydrogen bond donor to Leu455 backbone CO group within CDR-H1. In the context of CDR-H2, the hydroxyl group of Ser53HC targets both the backbone CO group of R457 and the hydroxyl group of Tyr421, the latter being involved in a hydrogen bond with the NH group of Gly54_HC_ as well; and the hydroxyl group of Ser56_HC_ interacts with the carboxyl group of Asp420. Overall, CDR-H2 interactions are less conserved among IGHV3-53/3-66 members compared to CDR-H1 interactions: the hydrogen bond between Tyr52_HC_ hydroxyl group and Lys417 amine group is observed only in B38 and CV30, while the hydroxyl group of Thr415 and the NH group of Gly416 are targeted by the hydroxyl group of Tyr/Phe58_HC_ only in B38, CV30 and 910-30. Structural comparison of light chain residues shows a strongly conserved tyrosine residue (Tyr32_LC_) in the CDR-L1 at the heavy:light chain interface which provides a stabilizing hydrophobic environment to the aromatic ring of Tyr505, together with Val28/29LC (in CV30, CC12.3, COVA2-04), Ile29_LC_ (in 910-30, B38, CB6, CC12.1) or Tyr30_LC_ (in C105). In the case of CC12.1, B38 and 910-30 Ser30LC interacts with the side chains of Gln498 and Asn501.

Using these defined sequence and structural motifs, we used published antibody repertoire data to estimate the prevalence of antibody class precursors in healthy human immune repertoires (Bräuninger et al., 2001; Sethna et al., 2019; Soto et al., 2019). Antibody lineages with anti-SARS-CoV-2 IGHV3-53/3-66 class characteristics were identified in approximately 1 in 44,000 reported human antibody sequences (**Fig. 3E**) (Soto et al., 2019), which we found to be a high frequency in comparison to previously studied anti-HIV-1 VRC01-class antibody precursors that occur in approximately 1 in 1-4 million human antibodies (Zhou et al., 2013). These comparatively high frequency estimates of IGHV3-53/3-66 anti-RBD precursors in human immune responses support the recovery of antibodies from this class in multiple convalescent COVID-19 patients.

### RBD up/down conformation influences S protein recognition for the IGHV3-53/3-66 antibody class

The RBD ‘up’ position is required for ACE2 engagement, as well as IGHV3-53/3-66 antibody binding (Du et al., 2020; Walls et al., 2020; Wrapp et al., 2020b). Cryo-EM structural analysis at endosomal pH has revealed a pH-mediated conformational switch that rotates RBD domains down at pH 5.5-4.5 (Zhou et al., 2020b). Because the recently emerged D614G mutation also alters the RBD ‘up’ vs. ‘down’ dynamics, we sought to understand how D614/D614G and pH-based alteration of ‘up’ vs. ‘down’ prevalence influence IGHV3-53/3-66 class recognition of spike (Walls et al., 2020; Zhou et al., 2020b). We investigated class binding at three pH values related to known RBD ‘up’ versus ‘down’ states for the D614 and D614G mutational variants (**Fig. 4A**) (Benton et al., 2020; Cai et al., 2020; Wrapp et al., 2020b; Yurkovetskiy et al., 2020; Zhou et al., 2020c). dhACE2 competition ELISA assays at pH 5.5 and 4.5 showed that IGHV3-53/3-66 class members compete in a concentration-dependent manner with dimeric human ACE2 for binding to SARS-CoV-2 S2P spike, and to the D614G S2P spike (**Fig. 4B, Supplemental Fig. 3**). Using single-cycle surface plasmon resonance, we found that the extremely potent mAb 1-20 recognized S protein and RBD with no loss in affinity at endosomal pH, whereas the less potent antibodies 910-30 and B38 showed reduced affinity in the endosomal pH range (**Fig. 4C, Supplemental Fig. 4**). We compared authentic virus neutralization IC50 potencies (from **Fig. 2D**) to the ratio of mAb-Spike affinity (**Supplemental Fig. 4**) divided by reported dhACE2-Spike affinity (Zhou et al., 2020b), which suggested that potent mAb neutralization was correlated with mAb affinity across all pH values tested (**Fig. 4D**). Finally, a qualitative Octet pH series analysis using D614 S2P spike showed that as the pH reduces (and RBDs preferentially rotate down), the potent neutralizer mAb 1-20 exhibited strong recognition of D614 S2P spike for pH≥6.0, whereas 910-30 showed reduced binding below pH=6.5, and the least potent B38 binding showed reduced binding below pH=7.0 (**Fig. 2E**, left panel). In contrast, all class members maintained strong binding to mutant D614G S2P spike into the endosomal pH range (where one RBD likely remains up), and the potent antibody class member 1-20 recognized D614G spike down to pH 4.0 (**Fig. 2E**, right panel). These data suggested that the most potent antibodies can maintain the bound state (and stabilize the RBD-up conformation) more effectively under endosomal pH conditions for D614 S2P spike, whereas all antibody class members could effectively recognize the native RBD-up conformation for D614G across a broad pH range. Our data support ACE2 competition as a functional signature of IGHV3-53/3-66 public antibody class neutralization, and we show that the RBD-up vs. RBD-down conformation substantially influenced the ability of IGHV3-53/3-66 class antibodies to recognize spike trimer.

**Fig. 4.**
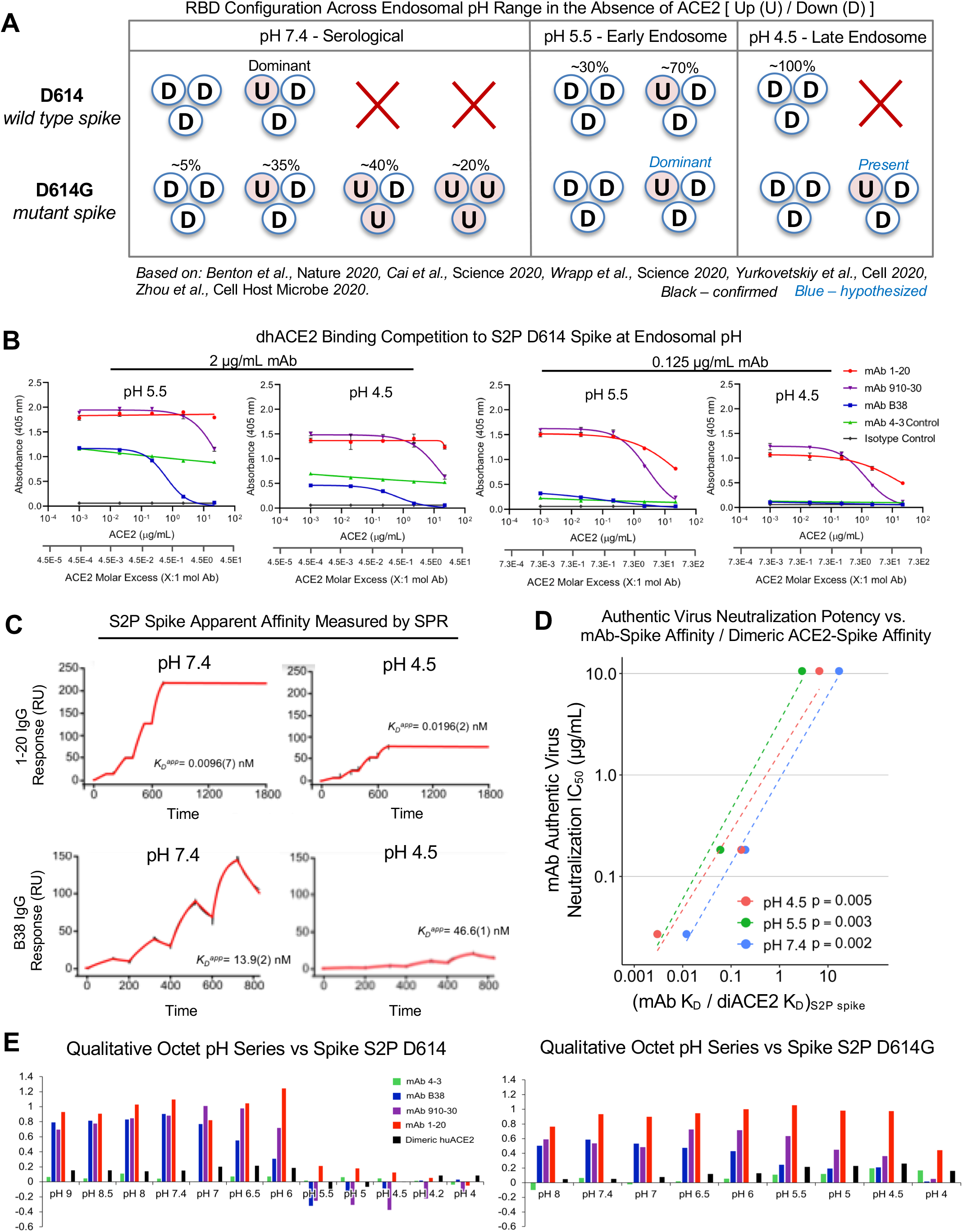
Up/down conformational changes of RBD influence IGHV3-53/3-56 antibody class recognition of spike protein across the serological to endosomal pH range. **(A)** Schematic of RBD conformational states inferred by Cryo-EM and experimental analysis for un-ligated D614 and D614G spike. U and D denote ‘up’ and ‘down’ RBD configurations, respectively. Percentages denote observed particle populations. **(B)** dhACE2 competition ELISA at endosomal pH against SARS-CoV-2 S2P protein. Constant IgG concentrations were used with varying dhACE2 (ACE2) concentrations at pH 5.5 and 4.5. dhACE2 concentration is provided as both μg/mL and as ACE2 molar excess units. **(C)** Single-cycle SPR kinetic assays for 1-20 and B38 IgG at serological and endosomal pH against biotinylated spike. Black traces represent experimental data and red traces represent the fit to a 1:1 interaction model. The number in parentheses represents the error of the fit in the last digit. **(D)** Correlations between authentic virus neutralization potency (from Fig. 2D) versus the ratio of antibody IgG affinity to spike S2P (from Fig. S4A) divided by dhACE2 affinity to spike S2P (reported from Zhou et al, 2020b). **(E)** Qualitative octet pH series for wild type S2P spike and escape mutant D614G S2P spike across a range of pH values. More potent IGHV3-53/3-66 class members retained binding at low pH against spike as compared to less potent class members.

## Discussion

Enhanced understanding of IGHV3-53/3-66 class-based spike recognition can provide insight into immune monitoring, antibody discovery, and vaccine design against SARS-CoV-2. Structural analysis of a novel class member mAb 910-30 revealed previously undescribed spike disassembly at high occupancy, and our antibody class comparative studies showed that native heavy:light pairing remains essential for potent neutralization, despite high similarities in heavy chain sequences. Comparative sequence-structure analyses enabled the identification of conserved light chain class signatures, defined as ^27a/28^[GDS]xSx{1,2}[FY] (kappa) and ^29^GY[KN] (lambda) residues in CDR-L1 that make key contributions to RBD recognition. We also note that class member light chains use common aromatic/hydrophobic residues ^28^Val, ^29^Ile/Val, or ^30/32^Tyr30/32 to achieve similar interactions with ^505^Tyr in the RBD, which is part of the shared ACE2 and IGHV3-53/3-66 class binding epitope. These shared light chain features illuminate the structural rationale for broader light chain diversity among IGHV3-53/3-66 class members.

The frequency of anti-SARS-CoV-2 IGHV3-53/3-66 precursor antibodies in healthy donors (around 1 in 44,000) was more common than the previously studied anti-HIV-1 VRC01-class antibody precursors observed in 1 per 1-4 million antibodies (Zhou et al., 2013). In addition, it has been shown that anti-HIV-1 VRC01-class antibodies also require much higher levels of somatic hypermutation (SHM) to achieve potent neutralization (Zhou et al., 2013). The comparably limited SHM required for anti-SARS-CoV-2 IGHV3-53/3-66 class antibodies appears to be a feature of IGHV germline gene neutralizing interactions and the need to recognize highly conserved viral variants, as compared to HIV-1 broadly neutralizing antibodies that must recognize broadly diverse viral variants and show limited germline gene neutralization. These findings help explain the observed reproducibility of public IGHV3-53/3-66 anti-RBD antibodies in convalescent COVID-19 patients.

D614G S2P spike variant shows a greater prevalence of RBD-up than D614G, which may enhance spike and the ACE2 host receptor recognition to confer higher D614G viral infectivity (Hou et al., 2020; Mansbach et al., 2020; Yurkovetskiy et al., 2020; Zhang et al., 2020). Conversely, a sustained RBD ‘up’ also could make the virus more sensitive to neutralization, as the exposed ‘up’ RBD enhances exposure of vulnerable epitopes (Mansbach et al., 2020; Zhou et al., 2020c). We outlined differences in RBD display caused by the D614G mutation that enhance antibody class recognition of spike across a broad pH range, and we show that D614G had no detrimental impact on IGHV3-53/3-66 antibody class neutralization, which agrees with prior reports (Plante et al., 2020; Weisblum et al., 2020; Weissman et al., 2020). Interestingly, only the most potent antibodies could bind to the D614 variant at endosomal pH which demonstrated that high-affinity antibody recognition can prevent D614 RBD from rotating down at pH 5.5-4.5. These data imply that screening for antibody recognition of D614 S2P at endosomal pH could be an effective method to identify potent anti-SARS-CoV-2 antibodies, and other studies have reported potent antibodies that recognize S trimer even in the context of RBD down conformations (Tortorici et al., 2020). We also found that ACE2 competition at pH 7.4 was correlated with potent antibody protection, consistent with the known cell surface attachment to ACE2 at serological pH.

In summary, here we report the discovery of a new public IGHV3-53/3-66 antibody class member and outlined the unique heavy and light chain interactions that lead to potent immune recognition of both D614G and D614G spike variants. These data enhance our understanding of the public IGHV3-53/3-66 antibody class and highlight its convergent neutralization features to accelerate anti-SARS-CoV-2 antibody mapping and inform future efforts to identify and elicit neutralizing antibody responses against COVID-19.

## Acknowledgments

We thank Jennifer Hackett from the Genome Sequencing Core Lab at the University of Kansas for help with Illumina sequencing, and R. Grassucci, Y.-C. Chi, and Z. Zhang from the Cryo-EM Center at Columbia University for assistance with cryo-EM data collection. Funding: This work was supported by the University of Kansas Departments of Pharmaceutical Chemistry and Chemical Engineering, COVID-19 Fast Grants, the Jack Ma Foundation, the American Lung Association, the Madison and Lila Self Graduate Fellowship Program, the Balsells Fellowship program, the Vaccine Research Center and the Division of Intramural Research of NIAID, NIH and by NIH grants DP5OD023118, R01AI141452, R21AI143407, and R21AI144408. This work was supported in part with federal funds from the Frederick National Laboratory for Cancer Research, NIH, under Contract HHSN261200800001.

## Author Contributions

B.B.B., G.C., A.S.F., C-H.S., S.N.L-A., K-T.Y, T.A.W., D.D.H., P.D.K., L.S., and B.J.D., designed the experiments; B.B.B., G.C., A.S.F., C-H.S., M.O., P.K., Y.T., P.W., M.S.N., Y.H., I.F., P.J.S., L.L., S.N.L-A., A.N., J.R.W., Y.L., X.P., B.M., A.D.L. and R.M. performed the experiments; A.S.O., I-T.T., J.Y., T.Z., E.R., and J.B. provided reagents for experiments, B.B.B., G.C., A.S.F., C-H.S., P.K., I.F., P.J.S., M.G-G., B.M., S.N.L-A., X.P, and B.J.D. analyzed the data; and B.B.B., G.C. A.S.F., C-H. S., P.D.K. L.S, and B.J.D. wrote the manuscript with feedback from all authors.

## Competing Interest Declaration

The authors declare no competing

Correspondence and requests for materials should be addressed to B.J.D.

## Supplementary Figure Titles and Legends

**Fig. S1.**
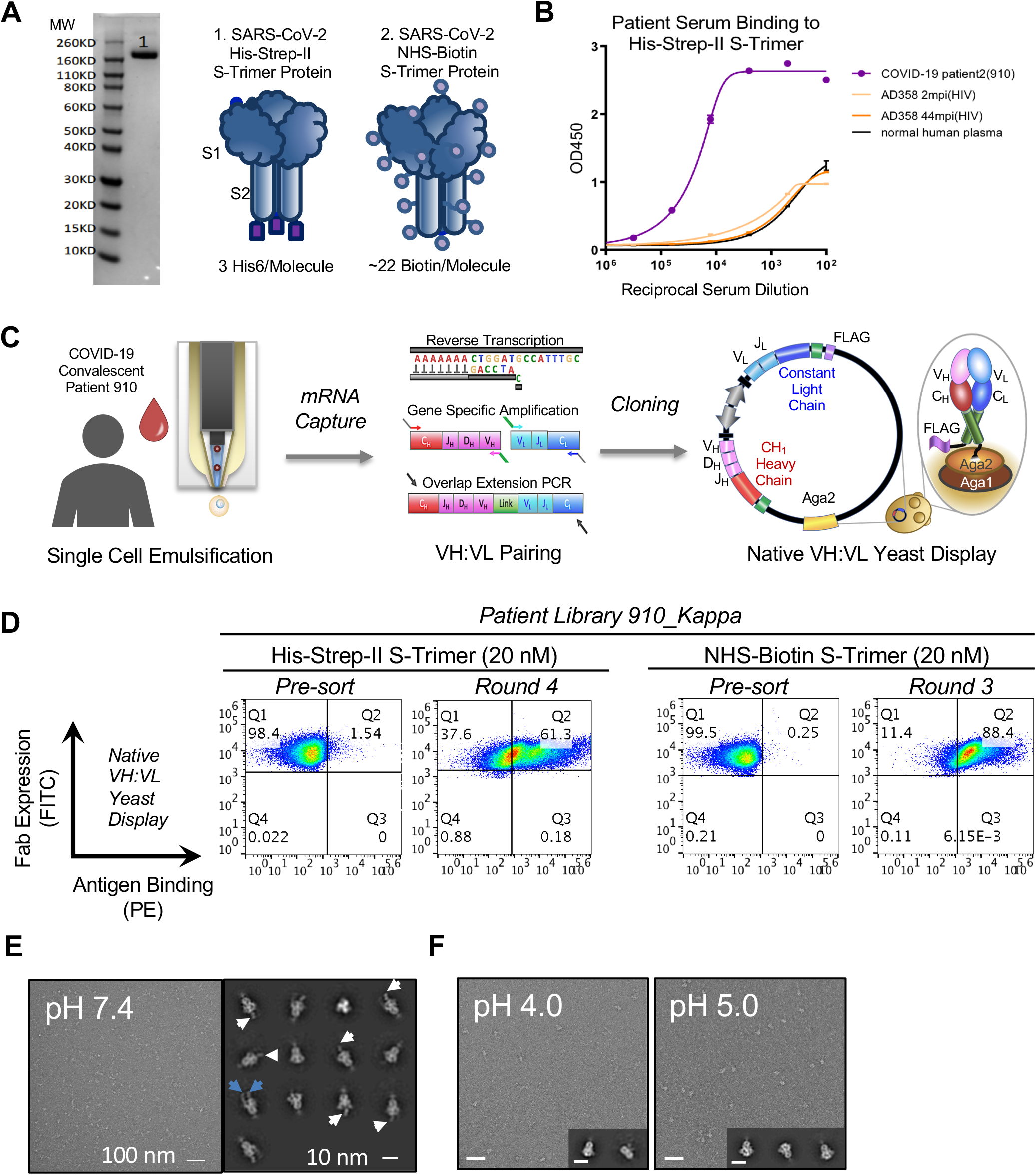
Overview of 910-30 discovery from a convalescent COVID-19 patient utilizing natively paired antibody fragment yeast display, FACS bio-panning, and soluble characterization. **(A)** Reduced SDS-PAGE gel shows SARS-CoV-2 His-Strep-II S-Trimer monomer protein at approximately 142 kDa. Schematics highlight unique features of SARS-CoV-2 Spike S2P antigen probes used for FACS biopanning of antibody yeast display libraries from a COVID-19 convalescent donor. **(B)** Hong Kong University convalescent Donor 910 serum showed strong binding to SARS-CoV-2 His-Strep-II S-Trimer protein compared to controls. **(C)** Workflow overview used to generate native VH:VL libraries from the COVID-19 convalescent donor HKU910 for functional antibody screening using yeast display. **(D)** Donor-derived antibody library bio-panning via FACS shows significant library enrichment after multiple rounds of sorting. *Left* Donor 910 pre-sort yeast library vs. sorted yeast library for His-S-Trimer antigen. *Right* Donor 910 pre-sort yeast library vs. sorted yeast library for Biotin-S-Trimer antigen. **(E)** Negative-staining electron microscopy at pH 7.4 resolved complexes between SARS-CoV-2 S2P and 910-30 Fab. Left: representative micrograph; right: representative 2D class averages. White arrows point to Fab fragments in complexes formed between one Fab and one spike trimer; blue arrows point to Fab fragments in complexes formed between two Fab fragments and one spike trimer. **(F)** Negative-staining electron microscopy at pH 4.0, and 5.0 reveal no 910-30 Fab bound to SARS-CoV-2 S2P protein at given pH values. Representative micrographs are shown. Insets show representative 2D class averages. Scale bars: 50 nm (micrographs), 20 nm (2D class averages).

**Fig. S2.**
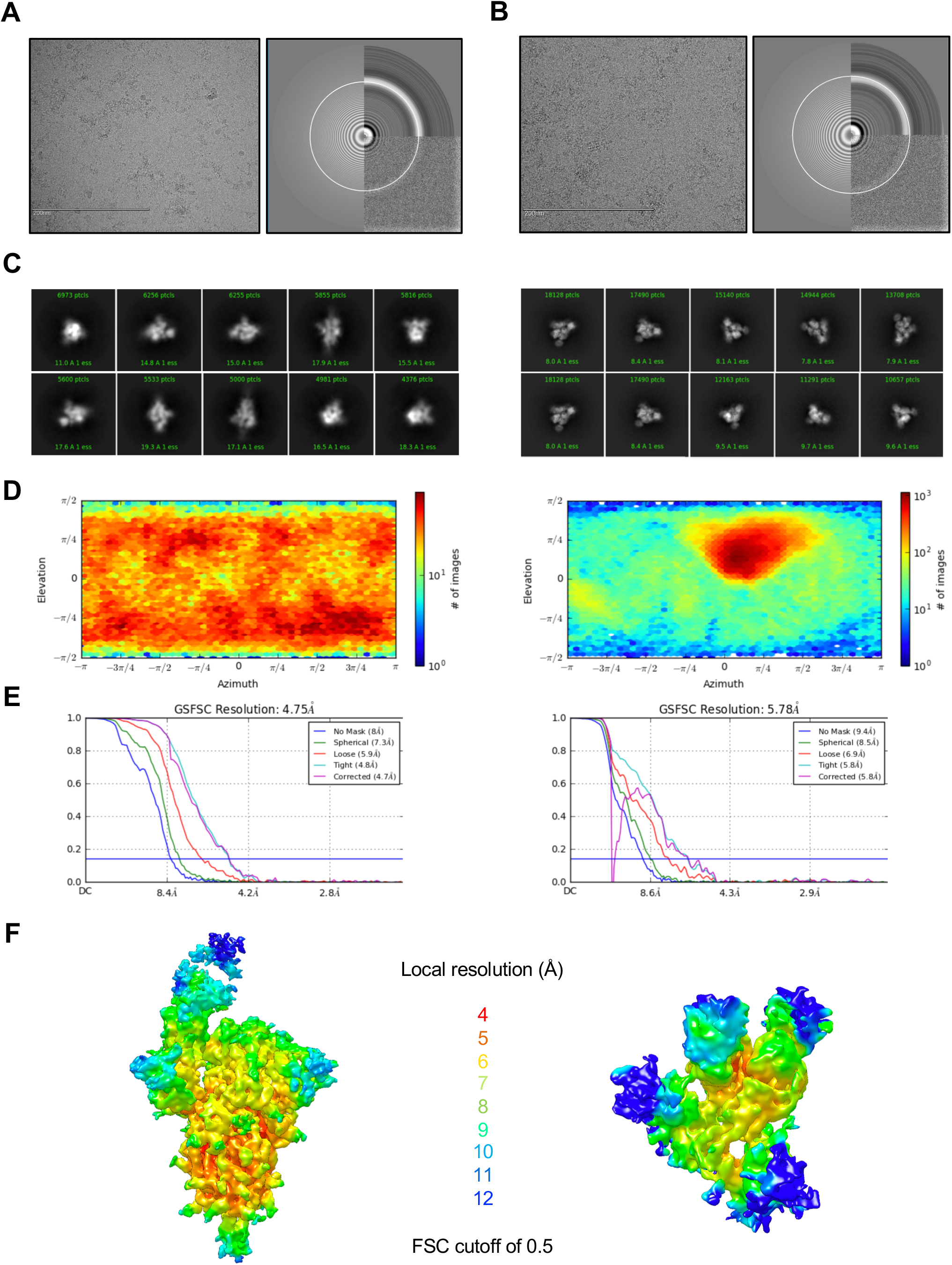
Cryo-EM analysis of 910-30 Fab in complex with SARS-CoV-2 spike at pH 5.5. Sample 1 obtained mixing 910-30 Fab and spike in a 1:1 molar ratio, sample 2 obtained mixing 910-30 Fab and spike in a 9:1 molar ratio. **(A)** Representative micrograph and CTF of the micrograph for sample 1. **(B)** Representative micrograph and CTF of the micrograph for sample 2. **(C)** Representative 2D classes for sample 1 (left) and sample 2 (right). **(D)** The orientations of all particles used in the final refinement are shown as a heatmap for sample 1 (left) and sample 2 (right). **(E)** The gold-standard Fourier shell correlation resulted in a resolution of 4.75 Å for sample 1 (left) and 5.78 Å for sample 2 (right). **(F)** The local resolution of the two final maps are shown generated through cryoSPARC using an FSC cutoff of 0.5; left: sample 1, right: sample 2.

**Fig. S3.**
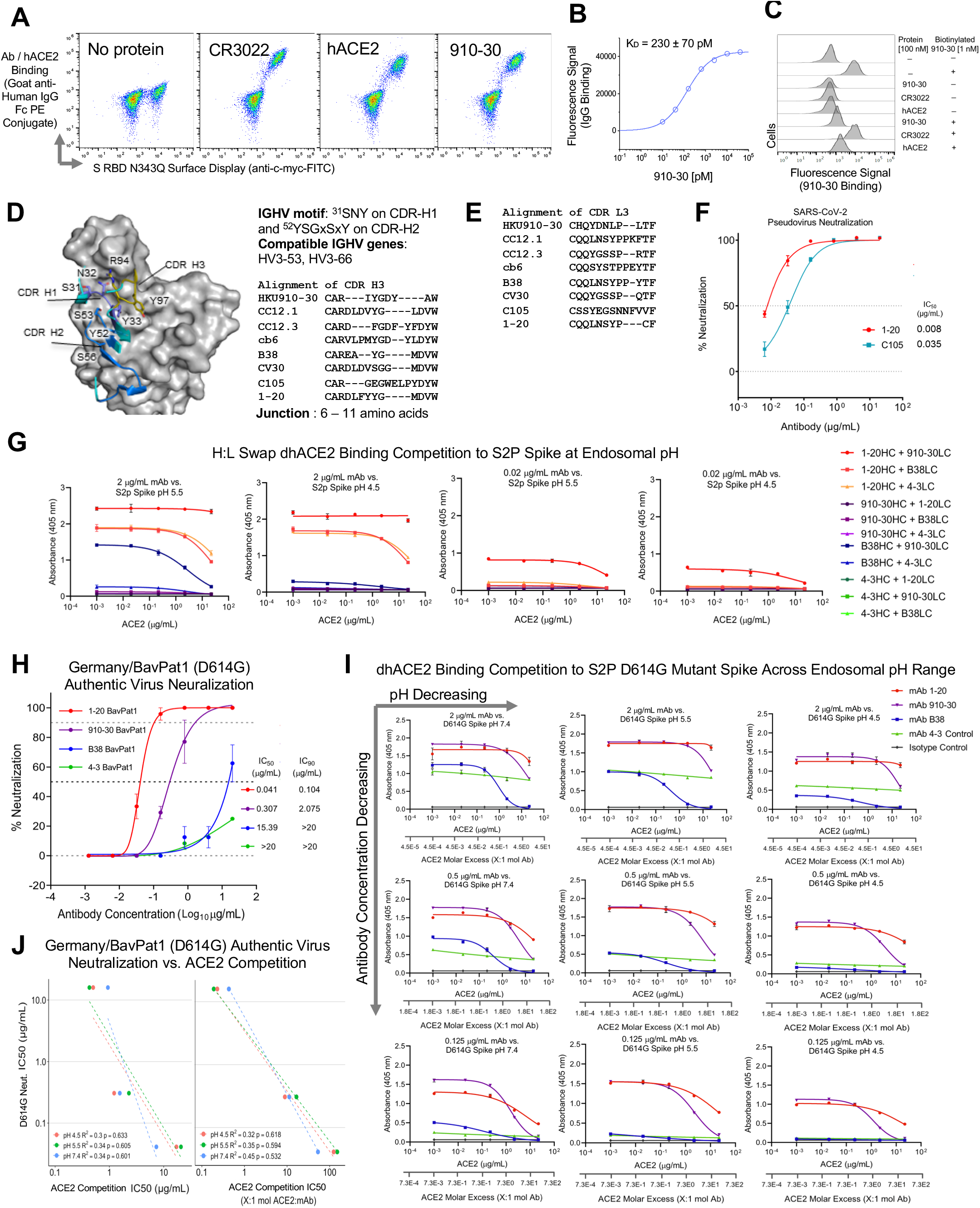
IGHV3-53/3-66 class member extended characterization and biophysical analysis. **(A)** Yeast-displayed aglycosylated RBD demonstrates 910-30 recognition is glycan-independent. S RBD N343Q with a C-terminal myc epitope tag was displayed on the surface of yeast and labeled with no protein or 1 nM of CR3022, human ACE2-Fc (hACE2), or 910-30. Cells were washed, secondarily labeled with anti-c-myc-FITC and Goat anti-Human IgG Fc PE conjugate, and read on a Sony SH800 cell sorter. Biological replicates were performed on two different days. **(B)** Yeast cell surface titrations of 910-30 IgG against aglycosylated S RBD yield a K_D_ of 230 ± 38 pM. Technical triplicates were performed for two biological replicates (n = 6), and error reported is 2 s.e.m. **(C)** Yeast-displayed RBD competition binding experiments of free 910-30, hACE2 and CR3022 vs. biotinylated or unbiotinylated 910-30. Technical triplicates were performed for two biological replicates (n = 6). **(D)** Heavy chain genetic elements associated with the IGHV3-53/3-66 antibody class. **(E)** Light chain CDR3 alignment of IGHV3-53/3-66 antibody class. **(F)** Lambda chain IGHV3-53/3-66 class member C105 shows moderate neutralizing capacity compared to potent kappa chain IGHV3-53/3-66 class member 1-20. **(G)** pH mediated dhACE2 competition measured by ELISA showing constant concentrations of heavy-light-swapped IgG binding to SARS-CoV-2 S2P protein versus increasing dhACE2 (ACE2) concentrations. Potently neutralizing heavy-light swap variants (Fig. 3A) show higher affinity binding to S2P spike and stronger ACE2 competition relative to less potent Abs. **(H)** 1-20, 910-30, and B38 show equivalent neutralization in a D614G authentic virus assay as for D614 authentic virus (Fig. 2D), with 4-3 included as a gene-matched control. **(I)** SARS-CoV-2 D614G S2P protein mutant variant pH mediated dhACE2 (ACE2) competition measured by ELISA showing constant concentrations of heavy-light-swapped IgG versus increasing dhACE2 concentrations. Potently neutralizing heavy-light swap variants show higher affinity binding to D614G S2P mutant spike and stronger ACE2 competition relative to less potent Abs. The concentration of dhACE2 required to outcompete antibody binding to spike is given as both μg/mL and as ACE2 molar excess units. **(J)** D614G authentic virus neutralization potency (Fig. S4H) and dhACE2 competition IC_50_ (Fig. S4I) show a correlation between potent neutralization and stronger ACE2 competition.

**Fig. S4.**
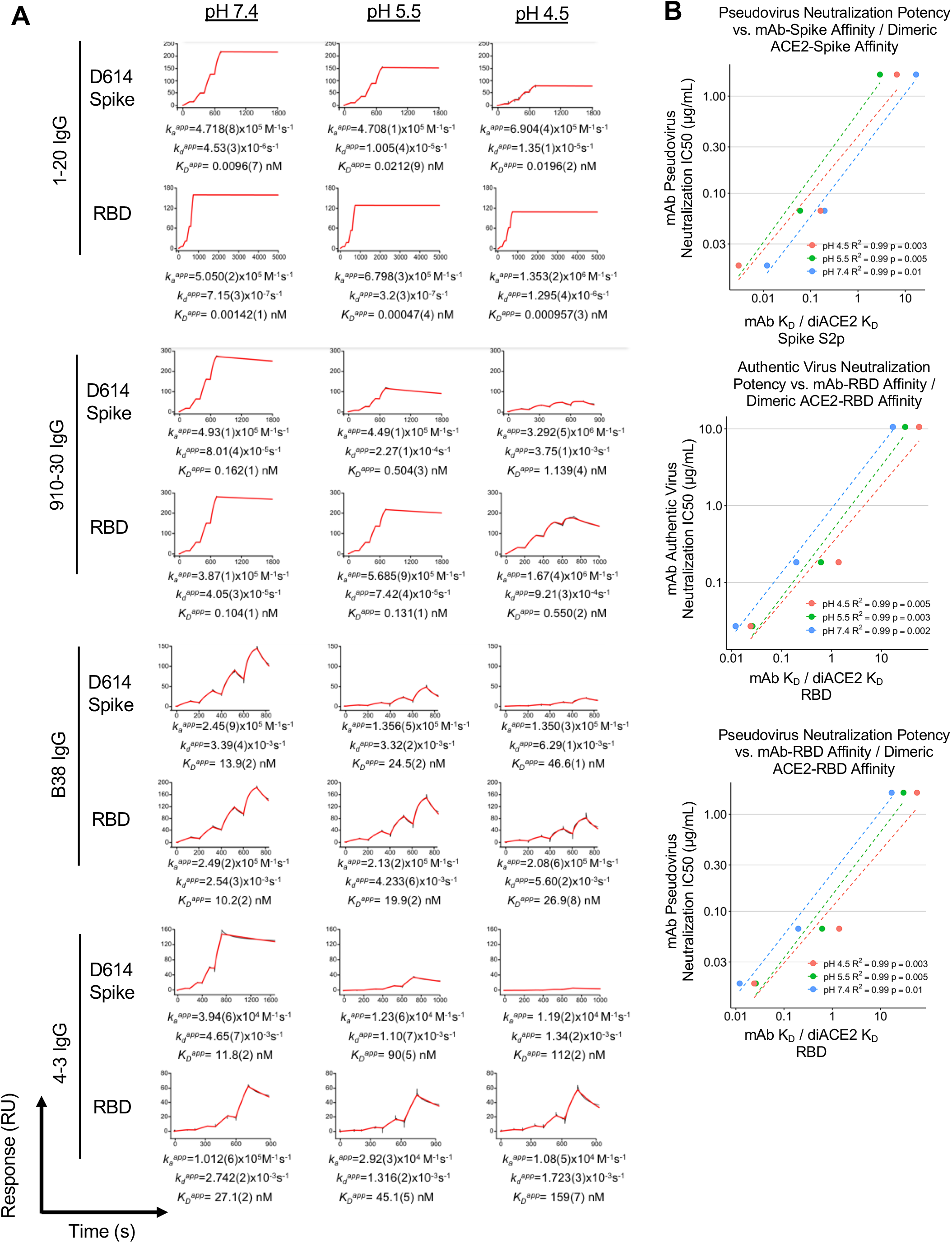
Extended binding and neutralization analysis across multiple pH values. **(A)** pH mediated SPR single-cycle kinetic experiments for 910-30, B38, 4-3, and 1-20, for IgG binding to biotinylated spike (top row) and to biotinylated-RBD (bottom row) in each of the four panels. Black traces represent the experimental data and red traces represent the fit to a 1:1 interaction model. The number in brackets represents the error of the fit in the last significant digit. **(B)** Correlations between both authentic and pseudovirus neutralization vs. the ratio of antibody IgG affinity to RBD or Spike divided by dimeric ACE2 affinity to RBD or Spike.

## Supplemental Table Legends

**Table S1.**
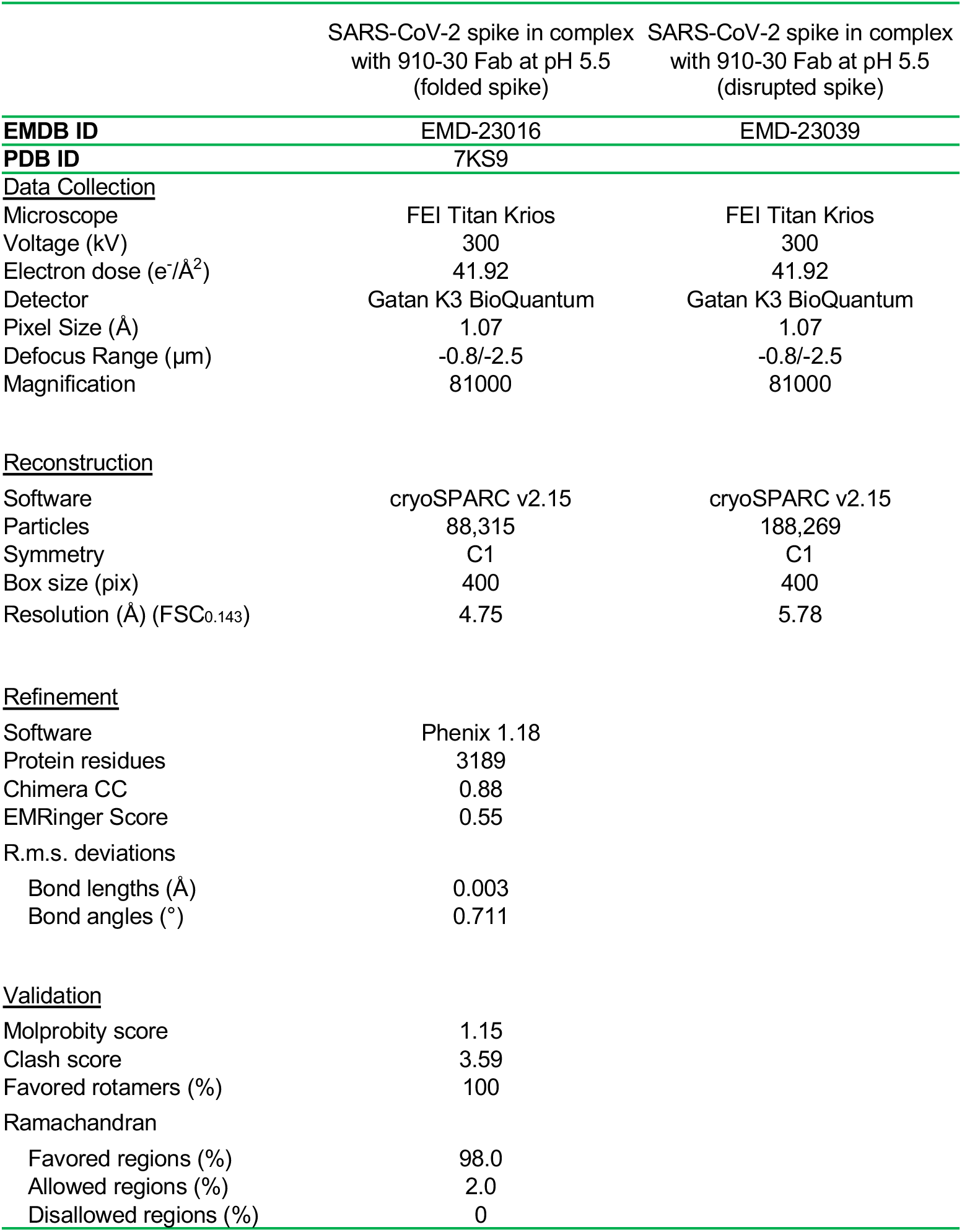
Cryo-EM data collection and refinement statistics for 910-30 Fab in complex with SARS-CoV-2 spike at pH 5.5.

**Table S2.**
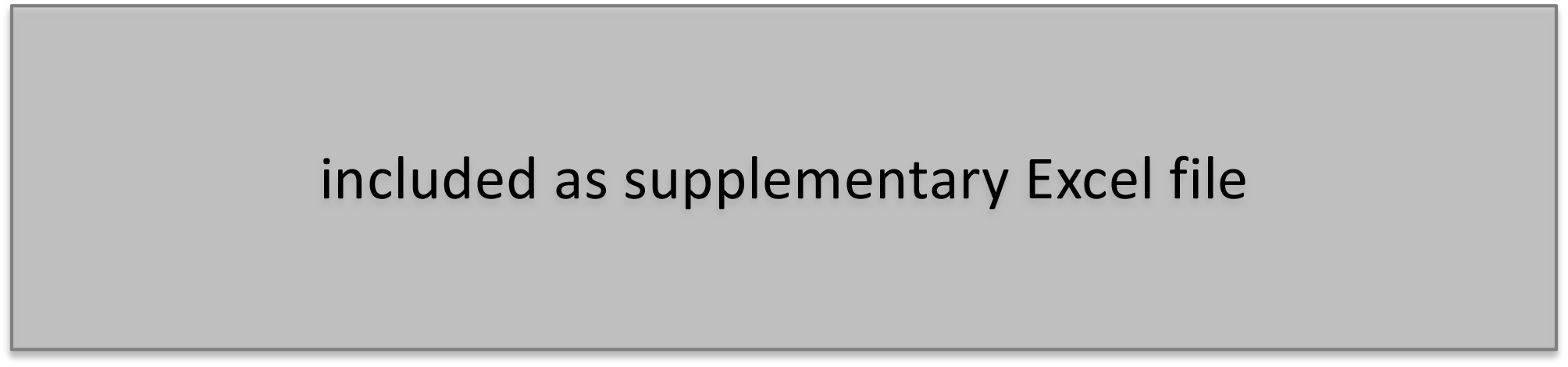
List of IGHV3-53 / IGHV3-66 anti-SARS-CoV-2 antibodies in previously published articles.

**Table S3.**
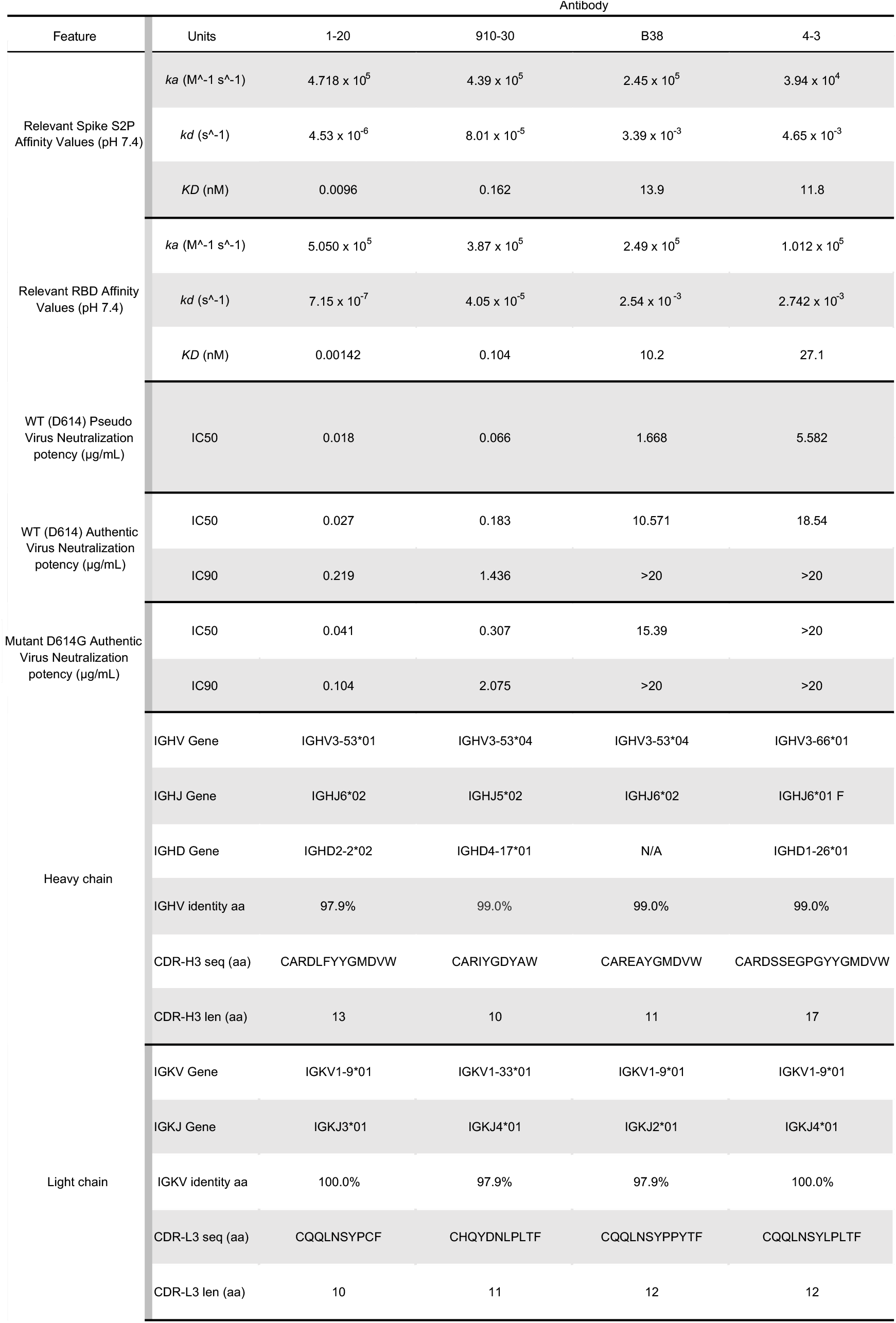
Features of the IGHV3-53/3-66 antibodies investigated in this study.

**Table S4.**
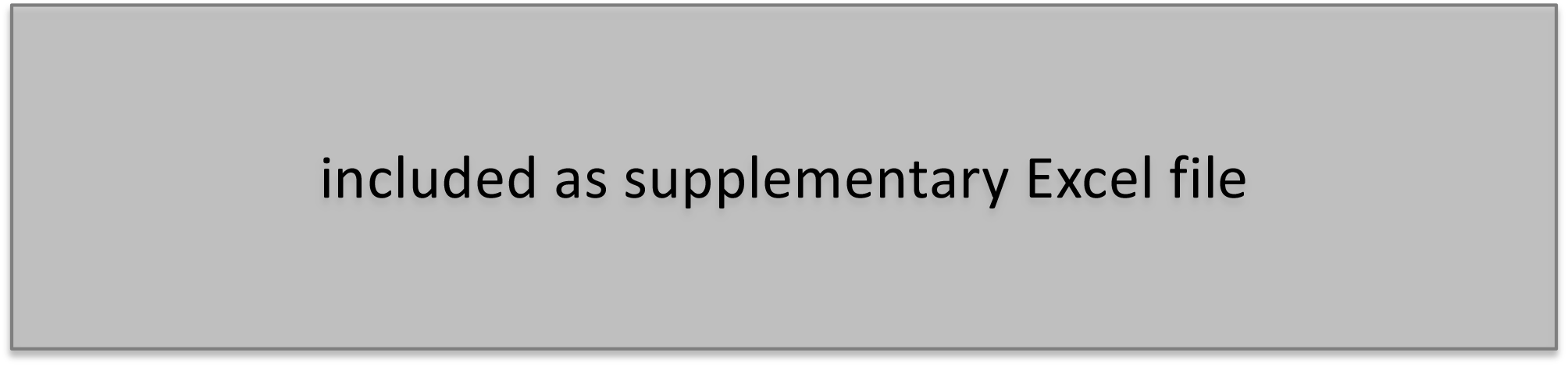
Heavy chain and light chain CDR1 and CDR 2 sequence alignment for recognition signature related to Figures 3C, S3D, and S3E.

## STAR Methods

### RESOURCE AVAILABILITY

#### Lead Contact

Further information and requests for resources and reagents should be directed to and will be fulfilled by the Lead Contact, Brandon J. DeKosky.

#### Materials Availability

Plasmids for antibody 910-30 generated in this study are available upon request for non-commercial research purposes.

#### Data and Code Availability

Cryo-EM coordinates and maps are deposited in the Protein Data Bank with accession code 7KS9 and in the Electron Microscopy Data Bank with accession code EMD-23016 (trimeric spike) and EMD-23039 (disrupted spike). The 910-30 neutralizing antibody variable heavy and variable light chain sequences have been deposited in GenBank with accession numbers MY291105 and MY291106, respectively. This study did not generate any unique datasets or code.

### EXPERIMENTAL MODEL AND SUBJECT DETAILS

Patient samples in the form of peripheral blood mononuclear cells (PBMCs) for B cell sorting were obtained from a convalescent SARS-CoV-2 patient, Donor 910. Cell line Expi293F cell was purchased from Thermo Fisher Scientific. Cell line HEK293T cell was purchased from Sino Biological. Cell line Vero C1008 (Vero-E6 cell) was purchased from ATCC. The cells were maintained and used following manufacturer suggestions and as described in detail below.

### METHOD DETAILS

#### Human Sample Collection

Informed consent was obtained for all study participants under IRB-AAAS9010 (Hong Kong University). Donor 910 (To et al., 2020) serum was collected for ELISA and neutralization assays, and PBMCs were cryopreserved for subsequent B cell receptor gene capture and antibody screening.

#### Expression and Purification of SARS-CoV-2 Antigens

The antigen probes used for sorting yeast surface displayed libraries were prepared as previously described (Liu et al., 2020a). Briefly, expression vectors encoding the ectodomain of the SARS-CoV-2 S protein was transiently transfected into Expi293 cells and then purified five days post transfection using on-column purification methods.

#### Production of SARS-CoV-2 Pseudovirus

SARS-CoV-2 pseudovirus was generated using recombinant Indiana vesicular stomatitis virus (rVSV) as previously described (Han et al., 2020; Liu et al., 2020a; Nie et al., 2020; Whitt, 2010). HEK293T cells were grown to 80% confluency then used for transfection of pCMV3-SARS-CoV-2-spike (kindly provided by Dr. Peihui Wang, Shandong University, China) using FuGENE 6 (Promega). Cells were cultured to grow overnight at 37 °C with 5% CO2. Then medium was removed and VSV-G pseudotyped ΔG-luciferase (G*ΔG-luciferase, Kerafast) was harvested to infect the cells in DMEM at a MOI of 3 for 1 h. Then cells were washed three times with 1 × DPBS. DMEM supplemented with anti-VSV-G antibody (I1, mouse hybridoma supernatant from CRL-2700; ATCC) and was added to the inoculated cells. The cells were then cultured overnight. The supernatant was removed the following day and clarified by centrifugation at 300 g for 10 mins before storing at −80 °C.

#### Emulsion Overlap Extension RT-PCR and Yeast Display Library Generation

B cells were isolated from Donor 910 cryopreserved PBMCs. Non-B cells were depleted by magnetic bead separation, and CD27^+^ antigen-experienced B cells were isolated by positive magnetic bead separation (EasySep Human B cell enrichment kit w/o CD43 depletion, STEMCELL Technologies, Vancouver, Canada, and CD27 Human Microbeads, Miltenyi Biotec, Auburn, CA, USA). Antigen-experienced B cells (memory B cells) were stimulated *in vitro* for 5 days to enhance antibody gene transcription. For stimulation, cells were incubated 5 days in the presence of Iscove’s Modified Dulbecco’s Medium (IMDM) (Thermo Fisher Scientific) supplemented with 10% FBS, 1x GlutaMAX, 1x non-essential amino acids, 1x sodium pyruvate and 1x penicillin/streptomycin (Life Technologies) along with 100 units/mL IL-2 and 50 ng/mL IL-21 (PeproTech, Rocky Hill, NJ, USA). B cells were co-cultured with irradiated 3T3-CD40L fibroblast cells that secrete CD40L (kind gift of John Mascola, Vaccine Research Center, NIAID) to aid B cell expansion. Single B cells were captured in emulsion droplets via a flow focusing device with concentric nozzles flowing suspended cells, lysis buffer with mRNA capture magnetic oligo (dT)-coated magnetic beads, and a viscous oil solution to form stable droplets compartmentalizing single B cells with lysis buffer and the mRNA capture beads (McDaniel et al., 2016). Captured beads loaded with single-cell mRNA were re-emulsified and the captured RNA product was reverse transcribed using a SuperScript™ III One-Step RT-PCR System with Platinum™ Taq DNA Polymerase (Thermo Fisher Scientific). The specific immunoglobulin VH and VL genes were then processed with an overlap-extension RT-PCR to link native heavy and light chains into a single amplicon, introducing two restriction sites: NheI and NcoI between the VH and VL genes for downstream yeast library generation (Wang et al., 2018). Natively paired antibody heavy and light chain sequencing and yeast surface display library generation were performed as described previously (DeKosky et al., 2013, 2015; Lagerman et al., 2019; McDaniel et al., 2016; Wang et al., 2018).

For yeast library generation, cDNA libraries were amplified with primers containing the yeast display vector restriction sites: AscI and NotI, used for subcloning into the yeast display vector. PCR amplified products were purified by agarose gel extraction and digested with AscI and NotI restriction enzymes followed by subsequent ligation into the yeast display vector backbone. This step was performed in duplicate for each library with separate Kappa- or Lambda-gene-specific primers and a corresponding Kappa or Lambda display vector to generate Kappa and Lambda libraries. Ligated plasmid libraries were transformed into high-efficiency electrocompetent E. coli, expanded overnight, and maxiprepped to isolate the plasmid library DNA product. Maxiprepped plasmid libraries were digested with NheI and NcoI restriction enzymes to remove the native linker from VH:VL pairing. Digested product was purified by agarose gel extraction, and then ligated with a pre-digested DNA gene encoding a bidirectional Gal1/Gal10 promoter inserted between the VH and VL sequences. The resulting ligated product was again transformed into high-efficiency electrocompetent E. coli, expanded overnight, and maxiprepped to isolate the plasmid library DNA product now containing the bidirectional promoter. A final PCR amplification was performed to amplify the VH:bidirectional promoter:VL amplicon with overhanging homologous ends to the pCT backbone for high-efficiency yeast transformation into AWY101 using an homologous recombination method previously described (Benatuil et al., 2010). Transformed libraries were passaged twice in SD-CAA to ensure a 1:1 ratio of plasmid DNA to yeast colony (Benatuil et al., 2010).

#### FACS Screening of Yeast Libraries

To induce Fab surface expression yeast libraries were incubated in SGD-CAA media at 20°C, 225 rpm for 36 hours. For the first round of sorting, 3×10^7^ presorted cells were washed twice with staining buffer (1x PBS with 0.5% BSA and 2 mM EDTA). Washed yeast display libraries were stained with 20nM of trimer antigen and a monoclonal anti-FLAG-FITC marker to measure Fab expression (Monoclonal ANTI-FLAG M2-FITC antibody, Sigma-Aldrich, St. Louis, MO, USA). For staining with the NHS-Biotin S-Trimer Protein probe, cells were mixed with 20nM un-labeled antigen and a monoclonal anti-FLAG-FITC marker (Monoclonal ANTI-FLAG M2-FITC antibody, Sigma-Aldrich, St. Louis, MO, USA) used to measure VL surface expression. This mix was incubated for 15 minutes at 4°C with gentle agitation on a platform shaker. Following incubation, a Streptavidin PE conjugate (Streptavidin, R-Phycoerythrin Conjugate Premium Grade, Thermo Fisher Scientific, Waltham, MA, USA) was added to the re-suspended mix to fluorescently label the biotinylated antigen protein and the sample was again incubated for 15 minutes at 4°C with gentle agitation on a platform shaker. These NHS-Biotin S-Trimer Protein samples were then washed 3x and re-suspended in a final volume of 1 mL in staining buffer before being filtered through a 35 micron-filter cap FACS tube. For staining the with His-Strep-II S-Trimer Protein probe, cells were incubated with un-labeled antigen for 15 minutes at 4°C with gentle agitation on a platform shaker. Samples were then washed 3x with staining buffer, and resuspended in a common mix containing the monoclonal anti-FLAG-FITC marker and a monoclonal anti-His-PE antibody (PE anti-His Tag Antibody, BioLegend, San Diego, CA, USA) to label surface expressed, antigen bound Fab. These cells were again incubated for 15 minutes in the fluorophore mix at 4°C with gentle agitation on a platform shaker. The fluorescently labeled samples were then washed 3x and resuspended in a final volume of 1 mL staining buffer before being filtered through a 35 micron-filter cap FACS tube. Samples were kept in the dark on ice until sorting. Subsequent rounds of enrichment sorting were performed using the same staining procedure, but for only 5×10^6^ input cells and 250 uL final resuspension volume.

A SONY Multi-Application 900 cell sorter running SONY LE-MA900FP Cell Sorter Software was used to detect all FITC+/PE+ cells from each sample and sort them into low pH SD-CAA media. The gating strategy used was previously described (Wang et al., 2018). Sorted yeast were expanded for 24-48 hours at 30 °C, 225 rpm and then passaged into SGD-CAA media to induce Fab expression for the next round of sorting. This process was repeated for 3-4 rounds of sorting to enrich for Fab-expressing antigen-binding library populations. In addition to the antigen-positive sorts, an aliquot of each yeast library was washed and stained with only the anti-FLAG-FITC marker, and all FITC+ (i.e., VL+) cells were sorted and sequenced for use as a reference database for NGS enrichment ratio calculations. Analysis of flow cytometry data was conducted using Flowjo10.4 (Flowjo, LLC, Oregon, USA).

#### NGS Analysis of Sorted Yeast Libraries

After each round of FACS enrichment, yeast libraries were expanded via incubation at 30°C for 24-48 hours. An aliquot of this culture was used for high-efficiency yeast plasmid DNA extraction (Whitehead et al., 2012). A high-fidelity polymerase (Kapa Hifi HotStart Mastermix, Kapa Biosystems, Massachusetts, USA) and primers targeting the yeast display vector backbone were used to amplify HC and LC genes from each library (Wang et al., 2018). A second round of primer-extension PCR with barcoded primers added a unique identifier to all HC and LC from a particular library (McDaniel et al., 2016). Sorted libraries were sequenced on the Illumina 2×300 MiSeq platform and sequencing was performed for each library after each round of FACS enrichment. Data processing of Illumina Raw FASTQ data was performed as reported previously (McDaniel et al., 2016; Wang et al., 2018). Briefly, Illumina sequences were quality-filtered to improve read quality, followed by V(D)J gene identification and annotation of CDR3 regions using IgBLAST (Ye et al., 2013). Antibody clonal lineages were tracked across yeast sort rounds by their CDR-H3 amino acid sequence and enrichment ratio. Enrichment ratios were calculated by comparing sequence prevalence in each sorted libraries to the unsorted, Fab-expressing (VL+) antibody library.

#### Antibody Production and Purification

The 910-30 antibody was codon-optimized, cloned into mammalian expression plasmid, and expressed as full human antibody IgG1s by co-transfection into Expi293 cells. Heavy and light chain plasmids were co-transfected into Expi293F (ThermoFisher) mammalian cells using the ExpiFectamineTM 293 Transfection Kit (Thermo Fisher Scientific, Massachusetts, USA) and culture in 37°C shaker at 125 RPM and 8% CO2. On day 6 post transfection, the supernatant from transient transfection culture were purified with Protein G or A resin (GenScript, New Jersey, USA) and concentrated using an Amicon Ultra-4 Centrifugal 30K Filter Unit (MilliporeSigma, Maryland, USA), then stored at 4°C.

#### ELISA Binding Assays to S trimer and RBD

S trimer and RBD enzyme-linked immunosorbent assays (ELISAs) (Fig. 2C) were performed in triplicate. 175 ng of antigen per well was coated onto 96-well ELISA plates at 4 °C overnight. Plates were washed and then blocked with 100 μL of blocking buffer at 37 °C for 2 hrs. Purified antibodies were serial diluted using dilution buffer, added to the antigen-coated blocked plates, and then incubated at 4 °C for 1 hr. Plates were washed and 50uL of a secondary anti-human kappa light chain detection antibody (A18853, Invitrogen, Carlsbad, CA) was added to each well and incubated at room temperature for 1 hr. After the final wash, 50 uL TMB substrate (00-4203-56, ThermoFisher Scientific, Waltham, MA) was used to detect antibody binding to antigen measuring absorbance at 405 nm.

#### Pseudovirus SARS-CoV-2 Viral Neutralization Assay

SARS-CoV-2 pseudovirus neutralization assays were performed as previously described (Liu et al., 2020a). Briefly, pseudovirus particles were generated from recombinant Indiana VSV (rVSV) expressing SARS-CoV-2 S protein. Neutralization was assessed by incubating pseudoviruses with serial dilutions of purified antibody, and scored by the reduction in luciferase gene expression.

#### Authentic SARS-CoV-2 Viral Neutralization Assay

Authentic virus neutralization assays were performed as previously described (Liu et al., 2020a). Briefly, to measure the neutralizing activity of purified mAbs an end-point dilution assay in a 96-well plate format was performed. Each mAb was 5-fold serially diluted starting at 20 μg/mL in triplicate. Dilutions were incubated with live SARS-CoV-2 for 1 hr at 37°C, and post-incubation the virus-antibody mixture was transferred onto a monolayer of Vero-E6 cells and incubated for 70 hrs. CPE from the resulting cell incubations were visually scored for each well in a blinded fashion by two independent observers.

#### dhACE2 Competition ELISA

Antibodies were assayed for dhACE2 competition by enzyme-linked immunosorbent assays (ELISAs) (Fig. 2E, 3C, S3I) in triplicate. ELISA experiments were performed in parallel at three pH values 7.4, 5.5, and 4.5. ELISA 96-well plates were coated with 175 ng per well of antigen in pH-adjusted PBS and incubated at room temperature for 1 hr. Ag-coated ELISA plates were washed and blocked with 100 μL of blocking buffer and incubated at room temperature for 1 hr. Purified antibodies were serial diluted and pre-mixed with dhACE2 using pH-adjusted dilution buffer. Ab:dhACE2 premixes were added to the pre-blocked, antigen-coated plates and incubated at room temperature for 2 hr. Plates were washed and 50 μL of 1:2000 diluted, pH-adjusted secondary anti-human kappa light chain detection antibody (A18853, Invitrogen, Carlsbad, CA) solution was added to each well and incubated at room temperature for 1 hr. After the final wash, 50 μL Super AquaBlue substrate was used to detect antibody binding to antigen measuring absorbance at 405 nm.

#### RBD Glycan Recognition Analysis via Yeast Display

For plasmid construction, pJS699 (S-RBD (333-537)-N343Q for fusion to the C-terminus of AGA2) was synthesized by PCR amplifying pUC19-S-ecto with primers PJS-P2196/PJS-P2197 (2.9kb) and PJS-P2198/PJS-P2199 (0.65kb). The resulting products were fractionated by agarose gel electrophoresis and the bands corresponding to the desired products were excised from the gel and purified using a Monarch DNA Gel Extraction Kit (NEB). The fragments were assembled using NEBuilder HiFi DNA assembly master mix (NEB) according to the manufacturer’s instructions and 5 μl of the reaction was transformed into chemically competent *E. coli* Mach1 (Invitrogen) and selected on LB agar supplemented with 50 μg/ml kanamycin.

To create the display construct of S-RBD (333-537)-N343Q fused to the C-terminus of Aga2p, pJS697 was digested with BsaI-HFv2 (NEB) and purified using a Monarch PCR & DNA Cleanup Kit (NEB). pJS699 was digested with NotI-HF (NEB), the reaction fractionated by agarose gel electrophoresis, and the band corresponding to S-RBD (0.83kb) excised and purified using a Monarch DNA Gel Extraction Kit (NEB). The two fragments were co-transformed (in a 2.4:1 molar ratio of S-RBD to backbone) into chemically competent *S. cerevisiae* EBY100 (Boder and Wittrup, 1997) and selected on M19D agar. M19D contained 5 g/L casamino acids, 40 g/L dextrose, 80 mM 2-(*N*-morpholino) ethanesulfonic acid (MES free acid), 50 mM citric acid, 50 mM phosphoric acid, 6.7 g/L Yeast Nitrogen Base (Sigma), and was adjusted to pH 7 with 9M NaOH, 1M KOH.

Recombinant human ACE2-Fc and CR3022 were received as a gift from Neil King and David Veesler at the University of Washington. Human ACE2-Fc was produced and purified as described (Walls et al., 2020). CR3022 (Ter Meulen et al., 2006) was expressed by transient transfection in Expi293F cells and purified by protein A affinity chromatography and SEC using a Superdex 200 10/300 GL. Specificity was verified by measuring binding to SARS-CoV-2 RBD and irrelevant antigen.

For yeast display screening, EBY100 harboring the RBD display plasmid was grown in 1 mL M19D overnight at 30°C. Expression was induced by resuspending the M19D culture to OD_600_=1 in M19G (5 g/L casamino acids, 40 g/L galactose, 80 mM MES free acid, 50 mM citric acid, 50 mM phosphoric acid, 6.7 g/L yeast nitrogen base, adjusted to pH7 with 9M NaOH, 1M KOH) and growing 22 h at 22 °C with shaking at 300 rpm. Yeast surface display titrations were performed as described (Chao et al., 2006) with an incubation time for 910-30 of 4 h and using secondary labels anti-c-myc-FITC (Miltenyi Biotec) and Goat anti-Human IgG Fc PE conjugate (Invitrogen Cat. No. 12-4998-82). Titrations were performed in biological replicate.

#### Glycosylation-Independent Binding for Antibody 910-30

EBY100 harboring the RBD display plasmid was grown in 1 ml M19D overnight at 30°C. Expression was induced by resuspending the M19D culture to OD600=1 in M19G (5 g/L casamino acids, 40 g/L galactose, 80 mM MES free acid, 50 mM citric acid, 50 mM phosphoric acid, 6.7 g/L yeast nitrogen base, adjusted to pH7 with 9M NaOH, 1M KOH) and growing 22 h at 22 °C with shaking at 300 rpm. Yeast surface display titrations were performed as described (Chao et al., 2006) with an incubation time for 910-30 of 4 h at room temperature and the secondary labels anti-c-myc-FITC (Miltenyi Biotec) and Goat anti-Human IgG Fc PE conjugate (Invitrogen Catalog # 12-4998-82). Titrations were performed in biological replicate (n = 2) with three technical replicates.

910-30 IgG was chemically biotinylated using NHS-Ester biotin (ThermoFisher EZ-Link Biotin Cat. No. 20217) at a 20:1 molar ratio of biotin:IgG according to manufacturer’s instructions. 1×10^5^ yeast cells were labelled with no protein or 100 nM non-biotinylated CR3022, hACE2 or 910-30 for 30 min at room temperature in PBSF (PBS containing 1 g/L BSA). The same cells were then labelled with 1 nM chemically biotinylated 910-30, in the same tube without washing, for 30min at room temperature in PBSF. The cells were centrifuged and washed with 200 μL PBSF. They were labeled with 0.6 μL FITC, 0.25 μL SAPE and 49.15 μL PBSF for 10 min at 4°C. Cells were then centrifuged, washed with PBSF, and analyzed on a flow cytometer. Experiments were performed with three technical replicates and two biological replicates.

#### Delineation of IGHV3-53/3-66 Sequence Signatures

A structure-based method was applied to define sequence signatures for the HV3-53/3-66 class COVID neutralizing antibody (Zhu et al., 2013). Briefly, protein structures of IGHV3-53/3-66 antibodies complexed with RBD or spike were selected for analysis, and the buried surface area (BSA) between antibody and RBD was calculated by the PDBePISA server (https://www.ebi.ac.uk/pdbe/pisa/). We examined the BSA larger than 20 Å^2^, and residues making contacting with the RBD projected surface that were encoded by the conserved germline sequence were selected as initial class sequence signatures, and amino acids from somatic hypermutations were used to refine the signature of the class antibody. For germline sequence alignments, heavy and light chain germline sequences were downloaded from IMGT (Lefranc et al., 2003) and the sequences of CDR1 and CDR2 were extracted and aligned based on Kabat numbering. ANARCI server was used to number amino acid sequences of antibody (http://opig.stats.ox.ac.uk/webapps/newsabdab/sabpred/anarci).

#### Antibody Class Frequency Estimation

The frequency of antibody class was estimated using OLGA software based on defined motif (Sethna et al., 2019). NGS samples of three healthy donors (NCBI Short Read Archive accession code: PRJNA511481) were used to analyze heavy and light chain lineage precursor frequencies (Soto et al., 2019). The ratio of human kappa and light chain (60:40) was obtained from Bräuninger et. al., 2001. Antibody class precursor frequency was calculated as:

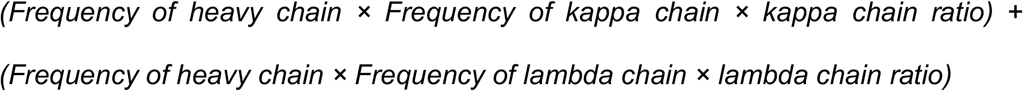

#### Negative Stain Cryo-EM

Samples were diluted to a spike concentration of about 20 μg/ml. A 4.7-μl drop of the diluted sample was applied to a glow-discharged carbon-coated copper grid. The grid was washed with a buffer with the same pH as the sample buffer (10 mM HEPES with 150 mM NaCl for pH 7.4; 10 mM acetate with 150 mM NaCl for the lower pH values). Protein molecules adsorbed to the carbon were negatively stained with 0.75% uranyl formate. Datasets were collected using a ThermoFisher Talos F200C electron microscope equipped with a Ceta CCD camera. The microscope was operated at 200 kV, the pixel size was 2.53 Å (nominal magnification: 57,000), and the defocus was set at −1.2 μm. Particles were picked and extracted automatically using in-house written software (YT, unpublished). 2D classification was performed using Relion 1.4 (Scheres, 2012).

#### Cryo-EM Sample Preparation

SARS-CoV-2 S2P spike was expressed and purified as described in Wrapp et al., 2020. 910-30 Fab was prepared by incubating the full 910-30 IgG with immobilized papain for 3 hours at 37 °C in 50 mM phosphate buffer, 120 mM NaCl, 30 mM cysteine, 1 mM EDTA, pH 7. Purified SARS-CoV-2 spike was diluted to a final trimer concentration of 0.33 mg/mL and mixed with 910-30 Fab in a 1:1 molar ratio (sample 1) or 1:9 molar ratio (sample 2). The final buffer for both samples was 10 mM sodium acetate, 150 mM NaCl, pH 5.5; 0.005% w/v n-Dodecyl β-D-maltoside (DDM) was added to the mixture to prevent aggregation during vitrification. After incubation for 1 hour on ice, a volume of 2 μL was applied to a glow-discharged carbon-coated copper grid (CF 1.2/1.3 300 mesh) and vitrified using a Vitrobot Mark IV with a wait time of 30 seconds and a blot time of 3 seconds.

#### Cryo-EM Data Collection, Processing and Model Fitting

Cryo-EM data were collected using the Leginon software (Suloway et al., 2005) installed on a Titan Krios electron microscope operating at 300 kV, equipped with a Gatan K3-BioQuantum direct detection device. The total dose was fractionated for 3 s over 60 raw frames. Data processing including motion correction, CTF estimation, particle picking and extraction, 2D classification, ab initio model generation, 3D refinements and local resolution estimation for both sample 1 and sample 2 datasets were carried out in cryoSPARC 2.15 (Punjani et al., 2017). The coordinates of SARS CoV-2 spike with 1 RBD up, PDB entry 6VYB (Walls et al., 2020), were employed as initial template to model the cryo-EM map of 910-30 Fab in complex with SARS-CoV-2 spike (sample 1). The RBDs were modeled using the crystallographic structure of RBD in complex with B38 Fab (PDB entry 7BZ5) (Wu et al., 2020b) as a template. The variable region of 910-30 Fab was initially modeled using PDB models 7BZ5 and 5SX4 (Sickmier et al., 2016) for the heavy and light chain respectively. The residues at the Fab:RBD interface were modeled by structural comparison of 910-30 Fab with 7 different antibodies belonging to the IGHV3-53/3-66 class. Automated and manual model building were iteratively performed using real space refinement in Phenix (Adams et al., 2004) and Coot (Emsley and Cowtan, 2004) respectively. EMRinger (Barad et al., 2015) and Molprobity (Davis et al., 2004) were used to validate geometry and check structure quality at each iteration step. UCSF Chimera (Pettersen et al., 2004) and Chimera X (Pettersen et al., 2020) were used to calculate map-fitting cross correlation (Fit-in-Map tool) and to prepare figures.

#### Octet Binding Experiments

Binding of mAbs 4-3, B38, 910-30, and 1-20 to SAR-CoV-2 S2P D614 and D614G variants was assessed an a FortéBio Octet HTX instrument (FortéBio). Experiments were run in tilted black 384-well plates (Geiger Bio-One) at 30°C and 1,000 rpm agitation. Running buffer was comprised of 10mM of the corresponding pH buffer plus 150mM NaCl, 0.02% Tween20, 0.1% BSA and 0.05% sodium azide. The following buffers were used to achieve the range of pH: pH 9 (borate), pH 8.5 (Tris), pH 8 (Tris), pH 7.4 (PBS), pH 7 (HEPES), pH 6.5 (MES), pH 6 (MES), pH 5.5 (NaAc), pH 5 (NaAc), pH 4.5 (NaAc), pH 4.2 (NaAc), pH 4.0 (NaAc). 300nM IgG solution was used for immobilization at pH 7.4 on anti-human IgG Fc capture biosensors (FortéBio) that were pre-hydrated for 30 minutes. Sensors were then equilibrated in 7.4pH buffer for 30 seconds followed by 180 seconds in the altered pH buffer. Binding was assessed at 200nM S2P D614 or D614G and response recorded for 180 seconds. Dissociation in the respective buffer was measured for 300 seconds. The Data Analysis Software HT v12.0 (Fortebio) was used to subtract reference well signal from loaded sensor dipped into buffer without spike protein. The maximum association response (nm) is reported at each pH.

#### SPR Binding Experiments

SPR binding experiments were performed using a Biacore T200 biosensor, equipped with a Series S SA chip. The running buffer varied depending on the pH of the binding reaction; experiments at pH 7.4 were performed in a running buffer of 10mM HEPES pH 7.4, 150mM NaCl, 0.1% (v/v) Tween-20; at pH 5.5 experiments were performed in 10mM sodium acetate pH 5.5, 150mM NaCl, 0.1% (v/v) Tween-20; and at pH 4.5 in 10mM sodium acetate pH 4.5, 150mM NaCl, 0. 1% (v/v) Tween-20. All measurements were performed at 25°C.

Biotinylated S2P was captured over independent flow cells at 750-900 RU. HKU910-30 and 1-20 IgGs were tested over the biotinylated S2P surfaces at four concentrations ranging from 1-27nM, while B38 and 4-3 were tested at four concentrations ranging from 3-81nM, to account for higher binding KDs. Biotinylated RBD was captured over independent flow cells at 250-500 RU and B38 was tested at four concentrations ranging from 3-81nM, HKU910-30 and 4-3 were tested at four concentrations of 1-27 nM and 1-20 at four concentrations ranging from 0.333-9 nM, to account for differences in their binding affinities. To avoid the need for surface regeneration that arises with the slowly dissociating interactions, we used single-cycle kinetics binding experiments. The four concentrations for each IgG were prepared in running buffers at each of pH, using a three-fold dilution series.

Binding of HKU910-30, 4-3 and B38 over the S2P or RBD surface as well as over a streptavidin reference surface was monitored for 120s, followed by a dissociation phase of 120s-1080s depending on the interaction at 50μL/min. For the interaction of 1-20 with the RBD, which showed an unusually slow dissociation rate, an extended dissociation phase of 4500s was necessary to extrapolate accurate apparent dissociation constants. Four blank buffer single cycles were performed by injecting running buffer instead of Fab to remove systematic noise from the binding signal. The data was processed and fit to 1:1 single cycle model using the Scrubber 2.0 (BioLogic Software). The results from these assays, are reported in terms of apparent kinetic parameters and K_D_s to account for potential avidity effects arising from the binding of bivalent IgGs to trivalent S2P.

### QUANTIFICATION AND STATISTICAL ANALYSIS

IC50 calculations were reported using GraphPad Prism software (version 8.4.3). Briefly, experimental data was imported and modeled using a least squares regression method to fit the data to a variable slope (four parameter) inhibitor vs. response curve with bottom parameters constrained to zero. Flow cytometry analysis was carried out using FlowJo software (version 10.4). The Spearman rank order correlation was calculated using cor.test function in base R. Spearman ρ and the p-values for the test were used to determine the strength of the correlation between tested variables.

### KEY RESOURCES TABLE

**Unique link for KRT:** https://star-methods.com/?rid=KRT5f99969f70be3

## Notes

### Competing Interest Statement

The authors have declared no competing interest.

